# Lysates of *Methylococcus capsulatus* Bath induce a lean-like microbiota, intestinal FoxP3^+^RORγt^+^IL-17^+^ Tregs and improve metabolism

**DOI:** 10.1101/855486

**Authors:** Benjamin. A. H. Jensen, Jacob B. Holm, Ida S. Larsen, Nicole von Burg, Stefanie Derer, Aymeric Rivollier, Anne Laure Agrinier, Karolina Sulek, Stine A. Indrelid, Yke J. Arnoldussen, Si B. Sonne, Even Fjære, Mads T. F. Damgaard, Simone I. Pærregaard, Inga L. Angell, Knut Rudi, André Marette, Jonas T. Treebak, Lise Madsen, Caroline Piercey Åkesson, William Agace, Christian Sina, Charlotte R. Kleiveland, Karsten Kristiansen, Tor E. Lea

## Abstract

Interactions between host and gut microbial communities may be modulated by diets and play pivotal roles in securing immunological homeostasis and health. Here we show that intake of feed based on whole-cell lysates of the non-commensal bacterium *Methylococcus capsulatus* Bath (McB) as protein source reversed high fat high sucrose-induced changes in the gut microbiota to a state resembling that of lean, low fat diet-fed mice, both under mild thermal stress (T_22°C_) and at thermoneutrality (T_30°C_). McB feeding selectively upregulated triple positive (Foxp3^+^RORγt^+^IL-17^+^) regulatory T cells in the small intestine and colon, and enhanced mucus production and glycosylation status suggesting improved gut health. Mice receiving McB lysates further exhibited improved glucose regulation, reduced body and liver fat along with diminished hepatic immune infiltration. Collectively, these data points towards profound whole-body effects elicited by the McB lysate suggesting that it may serve as a potent modulator of immunometabolic homeostasis.

## Introduction

Gut microbes shape intestinal immunity^1^ and increase the bioavailability of otherwise indigestible nutrients^2^. A well-balanced community structure is therefore essential for immunometabolic homeostasis, whereas aberrant gut microbiota compositions associate with numerous diseases, both within and outside the gastrointestinal tract^3^.

While therapeutic implications of rebalancing a mistuned gut microbiota appear promising, inconsistent response rates in relation to both probiotics and fecal transfer studies, with occasional adverse events, emphasize the complexity of such approaches. One example relates to the otherwise promising probiotic candidate *Akkermansia muciniphila*^4^, where negative effects have been seen in immunocompromised recipients^5, 6^. Similarly, *Prevotella copri* aids in metabolizing fibers in healthy individuals^7^ and protects against bacterial invasion in high fiber, chow-fed mice^8^, yet associates with insulin resistance in prediabetic obese individuals and precipitates glucoregulatory impairments in diet-induced obese (DIO) mice^9^.

An alternative to administering viable microbes is to utilize whole cell lysates, or selected cell components, of non-living bacteria. Apart from alleviating global energy demands, if used as a nutrient source, such components may also potently affect host physiology as recently reported for *A. muciniphila*^10^ and *Bifidobacterium bifidum*^11^. In the latter example, cell surface polysaccharides of *B. bifidum* were used to induce peripheral immune-tolerance via generation of regulatory T cells (T_regs_). The authors reported a pronounced increase in Foxp3^+^RORγt^+^ T_regs_ (_p_T_regs_), specifically in lamina propria (LP) of the large intestine (LI)^11^. This cell type is believed to be induced by commensal microbes and has emerged as a potent T_reg_ subset, exhibiting increased lineage stability and enhanced immunosuppressive capacity during intestinal inflammation compared to conventional Foxp3^+^RORγt^-^ T_regs_ (_n_T_regs_)^12^.

RORγt is the canonical transcription factor controlling IL-17 expression; a pleiotropic cytokine with both proinflammatory and immune-resolving actions depending on the eliciting cell type and physiological context^13^. Unfortunately, the studies describing _p_T_reg_ function did not measure IL-17 secretion. It therefore remains unknown whether these cells exhibit normal, reduced or increased IL-17 levels, and how this translates to host physiology. The impact of this predominantly colonic cell subset on host metabolism also remains unknown. Still, mounting evidence points towards the importance of intestinal IL-17 for controlling metabolic homeostasis^14, 15^. IL-17^+^ _p_T_regs_ may therefore be leveraged as a ‘dual hit’ strategy to curb immunometabolic dysfunction and gastrointestinal disturbances based on the immune-regulatory capacity of _p_T_regs_ concomitant with the metabolic benefits of gut-delivered IL-17.

To this end, we hypothesised that environmental bacteria, who have not been under evolutionary scrutiny for host-microbe interactions, would provide an unexplored reservoir of immunomodulatory stimuli. In support of this hypothesis, the methanotropic non-commensal bacterium *Methylococcus capsulatus* Bath (McB) has previously been shown to reduce inflammation and disease activity in dextran sulfate sodium (DSS)-induced colitis^24^. However, the impact on host metabolism and intestinal immune cells as well as mucus dynamics was not addressed.

We accordingly explored the effect of using whole-cell lysates from McB as protein source to reshape immunometabolism and the aberrant gut microbiota of DIO mice. We show that McB lysates augment Foxp3^+^RORγt^+^IL-17^+^ triple positive _p_T_regs_ in both SI- and LI-LP, and resets the obese microbiota concomitant with reversed key disease traits of diet-induced obesity.

## Materials and Methods

### Mice and Ethical Statements

All experiments were conducted in accordance with the EU directive 2010/63/EU as approved by the Danish Animal Experiments Inspectorate (#2014-15-2934-01,027). 6-7-week-old male C57BL/6JBomTac and C57BL/6JRj mice were acquired from Taconic Laboratories, Denmark, or Janvier Labs, France, respectively, as detailed below. All mice were allowed to acclimatize on regular chow diet for the first week upon arrival and then fed a LFD for two weeks prior to study initiation. Mice were kept under specific pathogen free conditions at 22 °C (T_22°C_) or 30 °C (T_30°C_), as indicated, in 12 h light/dark cycle (6AM–6PM).

### Housing, Diets and Experimental setup

Standard pelleted diets as well as protein-free western diet (WD) powder were obtained from Ssniff Spezialdiäten GmBH, Germany, and stored at -20°C throughout the duration of the experiment. Mice were fed a compositionally defined low fat diet (LFD, S8672-E050) until study initiation. A subgroup remained on LFD, whereas the remaining mice were transferred to a soy oil based reference WD (S8672-E025) (high fat, high sucrose, containing 0.15% cholesterol) for the run-in period (12 or 21 weeks dependent on the experiment). Experimental WDs were lot-matched and produced without protein (S9552-E021), which was subsequently added, and then pelleted, by the investigators as indicated; WD_REF_ = containing 19.5% (w/w) casein, WD_CNTL_ = containing 16.5% (w/w) casein and 3% Macadamia oil, WD_McB_ = 19.5% (w/w) whole-cell bacterial lysates (predominantly protein but up to ∼15% phospholipids^16^). Diet composition is more extensively described in Table S1. Detailed description of bacterial lysates is provided in the designated section below. All diets were freshly made for each experiment with independent batches of bacterial lysates to substantiate robustness and reproducibility of the observed findings. Mice were fed *ad libitum*, and feed intake was measured thrice a week. Water was changed weekly.

#### 12+6 protocol T_22°C_

Experimental setup is outlined in Figure 1A. C57BL/6JRj mice were purchased from Janvier Labs, France, and housed 3 mice per cage, reproduced in two independent experiments. For transparency, all data are depicted in respective graphs with mean ± SEM plus individual dots for the combined results of the two independent experiments. Experiment-dependent shapes enables visual discrimination of respective experiments as well as data distribution within and between experiments.

**Figure 1:**
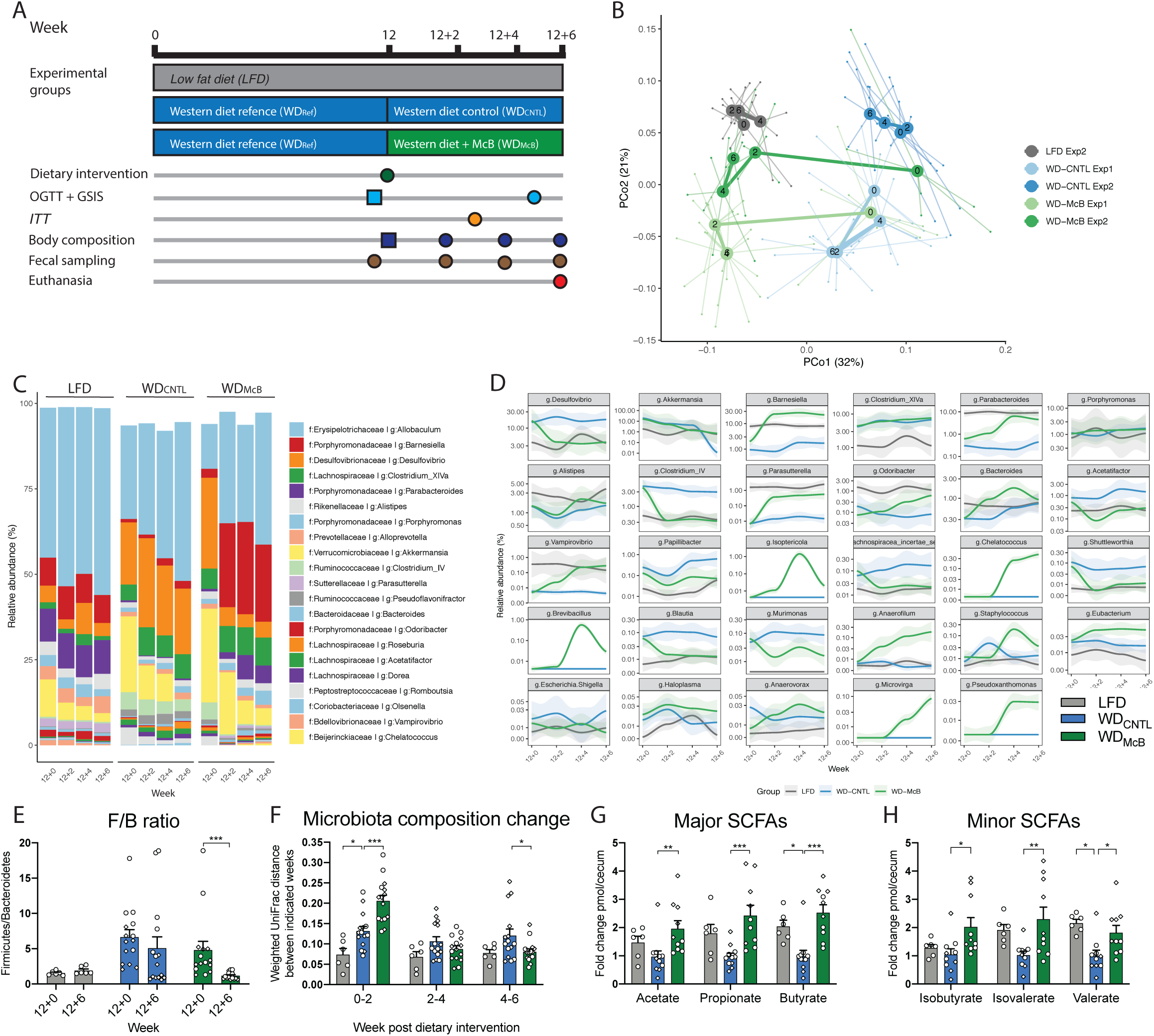
McB feeding reverses WD-induced gut microbiota changes and increases cecal SCFA levels: **A)** Study outline of two independent experiments with initial Western diet (WD_REF_) feeding for 12 weeks at 22°C and oral glucose tolerance test (OGTT), glucose-stimulated insulin secretion (GSIS), and body composition by MR scan assessed after 11 weeks to stratify groups (☐) followed by dietary intervention for 6 weeks of feeding either WD including McB lysate (WD_McB_) or a matched WD control diet (WD_CNTL_). During the experimental period body composition, OGTT, GSIS, and fresh fecal sampling was carried out at the indicated intervals. An insulin tolerance test (ITT) was carried out in the first of two experiment at the indicated timepoint. The second experiment included a reference group of fed low fat diet (LFD) fed mice. Groups or procedures featured uniquely in one experiment are italicized. **B)** Principle coordinate analysis (PCoA) using Weighted UniFrac distances of fecal microbiota sampled from first and second experiment biweekly during the dietary intervention period, as indicated by numbers post dietary intervention in centroids. The WD_CNTL_ and WD_McB_ groups were similar in microbiota composition prior to dietary intervention week 12+0 (PERMANOVA p = 0.88 and 0.43 in Exp1 and Exp2, respectively). At the end of each experiment, the microbiota composition was significantly different between these groups (PERMANOVA p = 0.001 and 0.002 in Exp1 and Exp2, respectively). **C)** Taxasummary of most abundant bacterial genera showing mean relative abundance in % of indicated family and genera in each group at indicated time points **D)** Deseq analysis of fecal bacterial genera abundances significantly regulated by McB intervention compared to the WD_CNTL_ (p.adj. < 0.05). Relative abundance in % in each group and variation are shown for each regulated genus at the sampled time points. Fold-change and adjusted p values of individual are indicated in Table S3. **E)** Firmicutes/Bacteroidetes ratio of fecal samples of individual mice before (12+0) and after 6 weeks of dietary intervention (12+6) in Exp1 (◆) and Exp2 (●). n = 6 (LFD) to 15 (WD_McB_ and WD_CNTL_). Statistical significance is indicated by * on paired t-test. **F)** Weighted UniFrac distance (instability test) between paired samples from indicated two-weeks’ interval post dietary intervention in Exp1 (◆) and Exp2 (●). n = 6 (LFD) to 15 (WD_McB_ and WD_CNTL_). Statistical significance is indicated by * on paired t-test. **G-H)** Major short-chained fatty acids (SCFAs) (acetate, propionate, and butyrate) and minor SCFAs (isobutyrate, isovalerate, and valerate) in cecal content as fold change pmol per cecum in Exp1 (◆) and Exp2 (●). n = 6 (LFD) and 10-11(WD_CNTL_ and WD_McB_, respectively). Statistical significance compared to WD_CNTL_ by one-way ANOVA with multiple comparison and Dunnett post hoc is indicated by *. **E-H)** * = p < 0.05; ** = p < 0.01, and *** = p < 0.001.

#### 21+5 protocol T_30°C_

Experimental setup is outlined in Figure 4A. C57BL/6JBomTac mice were purchased from Taconic Laboratories, Denmark, and initially housed 10 per cage to accelerate weight gain. 12 weeks into the run-in period, mice were single caged to allow accurate assessment of food intake, and also monitor if the more sensitive microbiome of single-housed mice would uniformly change towards a composition typical for LFD-fed mice, or if such phenomenon relied on a few highly responding mice transferring their microbiome to cage mates under group housed conditions. The timing of single housing allowed affected mice to adapt to social isolation and stabilize their weight development before intervention (Figure S4B).

### Cultivation of *M. capsulatus* Bath and preparation of bacterial lysate

*Methylococcus capsulatus* Bath (NCIMB 11132) was cultivated in nitrate mineral salts (NMS) medium as previously described^17^ to produce the single-strain bacterial lysate. McB culture aliquots were frozen in liquid nitrogen and stored at −80°C. Cultivations on agar plates and in shake flasks (orbital shaker incubator at 200 rpm) were performed at 45°C in an atmosphere of 75% air, 23.25% CH_4_, and 1.25% CO_2_. Continuous cultivation was carried out in a 3-liter bioreactor (Applikon, The Netherlands) with a working volume of 2 liters. Cells were pre-cultivated in shake flasks and used to inoculate the bioreactor to an optical density at 440 nm (OD_440_) of 0.1. The temperature was maintained at 45°C, stirring set to 650 rpm, and pH maintained at 6.8 by automatic addition of 2.5 M NaOH/2.5 M HCl. A gas mixture of 75% air and 25% methane was sparged into the bioreactor. The continuous culture was started after an initial batch phase, and the dilution rate was set to 0.01 h^−1^. The OD_440_ at steady state was generally sustained at approximately 10. Culture effluent was collected, and cells were harvested by centrifugation. Bacterial cell walls were disrupted by the use of a French press before freeze-drying of the material.

### Glucose and insulin tolerance tests

Mice were subjected to magnetic resonance (MR)-scan using EchoMRI 4in1 (Texas, USA) to determine fat- and lean mass at indicated time points (Figure 1A, 4A). Mice were fasted 5 or 2 hours prior to any oral glucose tolerance (OGTT) or intraperitoneal insulin tolerance test (ITT), respectively. Fasting blood glucose was measured by tail vein bleeding using the Bayer Contour glucometer (Bayer Health Care). Mice were subsequently gavaged with 3 µg glucose/g lean mass (OGTT) or intraperitoneally injected with 0.75 mU insulin/g lean mass (ITT). Blood were sampled at specified time points for blood glucose and insulin measurements to assess glucoregulatory capacity including glucose stimulated insulin secretion (GSIS) as described in detail elsewhere^18^.

### Short chain fatty acid measurements

Short chain fatty acids (SCFAs) were determined by gas chromatographic analysis. Feces were suspended in MilliQ water (1:1 (w/v)) and subsequently homogenized in a FastPrep-96 (MP Biomedicals) sample preparation unit. The homogenates were diluted with 0.4% formic acid (1:1 (v/v)), transferred to Eppendorf tubes and centrifuged at 13,000 rpm for 10 minutes. 300 μl of the supernatants were applied to spin columns (VWR, 0.2 μm pore size) and centrifuged at 10,000 rpm for 5 minutes. The eluates were transferred to 300 μl GC vials. Split injection mode was used with an injection volume of 0.2 μl. The gas chromatograph was a Trace 1310 (Thermo Scientific) equipped with an autosampler and a flame ionization detector. Helium was used as carrier gas, and the column was a 30 m long Stabilwax (Restek) with polyethylene glycol as stationary phase. Injector temperature: 250 °C, temperature intervals: 2 minutes at 90 °C before a 6 min increase to 150 °C, then a 2 minutes increase to 245 °C and a hold at this temperature for 4.9 minutes. Detector temperature: 275 °C.

### RNA extraction and quantitative RT-PCR

Frozen liver tissue was cryo-grinded by mortar and pestle on liquid nitrogen and total RNA was extracted by TRIreagent (Sigma-Aldrich, USA) according to the manufacturer’s protocol using PRECELLYS® 24 for homogenization. One µg of RNA was transcribed into cDNA by reverse transcriptase (Invitrogen, USA). Quantitative PCR analyzes were performed using the SYBR Green qPCR Master mix (Thermo Scientific, USA) and the Stratagene Mx3000P qPCR System. Standard curve and no template control (NTC) were included on every 96-well plates. Validation of each target was based on Rsq of standard curve >0.995, amplification efficiency of 100±10% and singular peak. Ct values for each target were related to 18S reference Ct values of the same sample by 2^ΔCt^ and visualized as fold-change to the LFD reference group mean. Primer sequences are summarized in Table S2.

### Histology

Liver tissue was sampled promptly after euthanasia, fixed in 10% formalin and subsequently embedded in paraffin prior to sectioning according to standard procedures for light microscopy. Colon was emptied for content and subsequently rolled around a 27G needle after which the generated Swizz Rolls were carefully preserved in liquid nitrogen (wrapped in aluminum foil) until paraffin embedding. Embedded tissues were cut 2 µm thick, and sections mounted onto glass slides. These sections were processed further and stained either with hematoxylin and eosin (H&E), Oil Red O, PAS, or HID-AB, or labelled with antibody of interest for immunohistochemical investigations, as detailed below. Samples were randomized and blinded to the pathologists performing the histological analyses.

#### Liver

##### Non-alcoholic fatty liver disease activity score (NAS)

Liver sections were prepared by standard protocols and subsequently assessed by two independent observers in a blinded fashion, using the established NAFLD activity score (NAS) for evaluation of H&E stained liver sections^19^. In short, score of 0 to 2 excludes non-alcoholic steatohepatitis (NASH), a score of 3–4 defines “borderline NASH”, and score ≥ 5 is considered as NASH.

##### Oil Red O staining

Liver biopsies were fixed in paraformaldehyde, cryoprotected in polyvinylpyrrolidone/saccharose, and frozen in liquid nitrogen. Cryosections were prepared and stained using the lipid-specific Oil red O (Sigma Aldrich, Germany) and analyzed under an Axio Imager M2 microscope. Images were captured with the Axiocam 506 color camera (Zeiss, Jena, Germany). Areas of stained lipids were determined using ImageJ-1.50i software (https://imagej.nih.gov/ij/index.html).

#### Colon

The morphological and morphometrical evaluations of the colonic tissues were performed in a blinded fashion, and all measurements were made by the same trained pathologist. One H&E stained section of colon from each individual was evaluated histologically for any sign of pathology. To assess the quality of the mucins, colonic tissues were stained for acidic as well as neutral mucins. Neutral mucins were stained by Periodic Acid Schiffs (PAS). Acidic mucins were stained through a combination of High Iron Diamine (HID), for sulfated mucins (sulfomucins), and Alcian Blue (AB) for carboxylated mucins (sialomucins). Crypt depth (CD) measurements, as well as analyses of mucin type and amount, were performed on HID/AB- and PAS-stained sections in three histologically distinct areas of the colon, i.e. the proximal, middle and distal area. At each of these three locations, three longitudinally sectioned crypts were measured. Each data point is thus presented as an average of three (mucin type and amount) or six (CD) measures. Crypts were selected only when the entire crypt epithelium was visible from the *lamina muscularis mucosae* to the lumen. The histological sections of the colon were examined in an Axio Imager Z2 microscope (Zeiss, Jena, Germany), and digital images were obtained using an Axiocam 506 color camera (Zeiss, Jena, Germany). Micrographs were captured with the same 20× objective magnification. Crypt depth measurements were performed using the software program Image J-Fiji version 1.52e Java 1.8.0_181^20^. For quantitative evaluation and automated scoring of the different mucin types, a plugin for color deconvolution for the Image J-Fiji program was used^21^.

### Hepatic lipidomics

Samples were randomized and lipid extracted as previously described. In short, 50 mg of frozen tissue were homogenized in a pre-chilled 2 mL tube added 0.5 mL of ice-cold 50% MeOH with ∼40 µM D5-tryptophan following standard protocols and stored at -80°C until LCMS analysis. The peaklist obtained after preprocessing (features defined by their mass/charge value, retention time and peak area) was analyzed in MetaboAnalyst 4. Data were then auto-scaled (mean-centering and division by the square root of standard deviation of each variable) to enforce Gaussian distribution enabling relative comparison. Univariate and multivariate analysis were performed with t-tests, volcano plot, analyses of variance (ANOVA) followed by Tukey’s HSD test, and principal component analysis (PCA) to detect significant hits and to visually separate trends between groups. Features showing a similar pattern between LFD and WD_McB_ were identified by pattern matching function. The top 25 features obtained were then processed for identification using the online database Lipidmaps (http://lipidmaps.org) with a mass tolerance between the measured m/z value and the exact mass of 3 ppm.

### Microbiome analysis and bioinformatic processing

Fresh feces samples were collected 3-4 hours into the light cycle, snap frozen and stored at -80°C until downstream processing. Bacterial DNA from fecal samples was extracted using a NucleoSpin soil kit (Macherey-Nagel) according to manufacturer’s instructions. 16S rRNA gene amplification, library preparation and sequencing were performed as previously described^22^. Initial data processing was performed as described elsewhere^23^. Data was rarefied to 23,107 reads per sample. The principal coordinate analysis (PCoA) was conducted based on the weighted UniFrac distance. To select OTUs and species enriched in different subgroups, DESeq2^24^ with default settings was used. The relative abundances of OTUs were aggregated to genus level. Genera present in >1/3 of the samples were used in the differential abundance testing. Genera with p.adj. < 0.05 comparing the last sampling point for WD_REF_/WD_CNTL_ vs WD_McB_ with DESeq2 differential expression analysis based on the negative binomial distribution were reported. Permutational multivariate analysis of variance (permanova) using weighted UniFrac distance matrices (Adonis, Vegan R package) was used to evaluate overall microbial compositions.

### Isolation of small intestine (SI) and large intestine (LI) lamina propria (LP) cells

Feces and mucus from LI were scraped off while SI was flushed with 1x HBSS (Gibco) containing 15 mM HEPES (Thermo Scientific). Peyeŕs patches were carefully excised from SI. SI and LI were opened longitudinally, cut into approx. 1 cm pieces and washed 3 times with 1x HBSS (Gibco) containing 15 mM HEPES (Thermo Scientific), 5% FCS (Viralex^TM^, PAA Laboratories), 50 µg/mL gentamycin (Gibco) and 2 mM EDTA (Invitrogen, Life Technologies) for 15 min at 37°C. During the first incubation step, 0.15 mg/mL DL-dithiothreitol (Sigma-Aldrich) was additionally added to LI samples. LI, but not SI samples, were shaken on an orbital shaker at 350 rpm during incubation. After each incubation step, SI, but not LI samples, were shaken by hand for 10 s. Media containing epithelial cells and debris were discarded by filtration through a 250 µm mesh (Tekniska Precisionsfilter, JR AB). The remaining tissue was digested for 25-28 min at 37 °C under magnetic stirring (500 rpm) in R-10 media (RPMI 1640 (Gibco) with 10% FCS (Viralex, PAA Laboratories), 10 mM HEPES (Thermo Scientific), 100 U/mL penicillin (Gibco), 100 µg/mL streptomycin (Gibco), 50 µg/mL gentamycin (Gibco), 50 µM 2-mercaptoethanol (Gibco) and 1 mM sodium pyruvate (Gibco)) containing 1 mg/ml collagenase P (Roche) and 0.03 mg/mL DNase I (Roche). After digestion, SI-LP and LI-LP cells were purified by density gradient centrifugation (600 *rcf* for 20 min at 22 °C, acceleration 5 and brake 0) with 40/70 % Percoll (GE Healthcare). Cell suspensions were subsequently filtered through 100 µm cell strainers (BD Biosciences) and re-stimulated *in vitro* prior to staining for flow cytometric analysis.

### *Ex vivo* stimulation of SI-LP and LI-LP cells

SI-LP and LI-LP cells were re-stimulated *in vitro* as previously described^25^. Briefly, cells were simulated in R-10 medium in the presence of either 20 ng/mL IL-23 (R&D Systems) or 250 ng/mL PMA (Sigma-Aldrich) in combination with 0.5 µg/mL ionomycin (Sigma-Aldrich) for 4 hrs at 37°C. After 1 hr incubation, 10 µg/mL brefeldin A (BioLegend) was added.

### Flow cytometry

Flow cytometry was performed according to standard procedures. Cell aggregates (identified in FSC-H vs. FSC-A scatter plots) and dead cells identified by using Zombie Aqua Fixable Viability Kit (BioLegend) were excluded from analyses. Intracellular staining was performed using the eBioscience FoxP3/Transcription Factor Staining Buffer Set (eBioscience) according to the manufacturer’s instructions. Data was acquired on a LSRII (BD Biosciences) and analyzed using FlowJo software (Tree Star).

### Antibodies (Abs) and reagents

The following mAbs and reagents were used in the study: anti-CD3ε (17A2), anti-CD4 (GK1.5), anti-CD8α (53-6.7), anti-CD11c (N418), anti-CD19 (6D5), anti-CD45 (30F11), anti-CD45R/B220 (RA3-6B2), anti-CD90.2 (30-H12), anti-CD127 (A7R34), anti-Ly-6G/Ly-6C (Gr-1) (RB6-8C5), anti-Ter119 (TER-119), anti-TCRβ (H57-597), anti-TCRγδ (GL3), anti-NK1.1 (PK136), anti-NKp46 (29A1.4), anti-FoxP3 (FJK-16s), anti-RORγt (B2D), anti-T-bet (4B10), anti-IL-17A (TC11-18H10.1), anti-IL-22 (IL22JOP) and anti-IFNγ (XMG1.2). All Abs were purchased from eBioscience, BioLegend or BD Biosciences. PECF594-conjugated streptavidin was purchased from BD Biosciences.

### Multiplex cytokine quantification

Liver tissues were crushed with a mortar and pestle on liquid nitrogen. 30-40 mg of tissue were transferred to a clean tube and added 400 µL T-PER tissue protein extraction reagent (Thermo Scientific) buffer including proteinase inhibitors (Sigma-Aldrich). Protein concentration was assessed by BCA (Thermo Scientific) according to manufacturer’s instruction. Cytokines were quantified using xMAP technology with combinations of Bio-Plex Pro (BioRad) premixed mouse cytokine panels, and the analyses were carried out on a Bio-Plex 200 system and the Bio-Plex Manager Software package.

### Statistical analysis

Omics data were acquired as described in their appropriate sections. Normality residuals of remaining data were assessed by D’Augostino-Pearson omnibus (k2), Graphpad Prism 8, and subsequently evaluated by parametric (gaussian distributed) or non-parametric (non-gaussian distributed) tests as appropriate. Details of each test including post hoc assessment are specified in figure legends. Data are expressed as mean ± SEM with individual dots, and all groups are compared to the relevant WD_REF_ or WD_CNTL_ group as indicated.

## Results

### McB feeding reverses WD-induced gut microbiota changes and increases cecal SCFA levels

To induce obesity and immunometabolic dysfunctions, C57BL/6JRj mice were initially fed an obesogenic WD. After 12 weeks of WD feeding, the mice were stratified into new groups based on weight, fat mass and glucoregulatory capacity (Figure 1A), and fed experimental WDs for an additional 6 weeks. While dietary fat is known to elicit reproducible and lipid-dependent alterations in the murine gut microbiome across a variety of different diet compositions^26^, less is known about the microbiome-modulating impact of protein. Thus, to investigate if dietary protein (i.e. casein versus whole-cell bacterial lysates) would affect gut microbiota community structures, we analyzed freshly collected fecal samples before and throughout the dietary intervention. LFD and WD_REF_ fed mice showed distinct gut microbiota profiles after 12 weeks of feeding (intervention baseline, week 12+0; Figure 1B,C), including ∼10-fold lower abundance of the health-promoting genera, *Parasutterella*^27^ and *Parabacteroides*^28^, countered by an equally increased abundance of the obesity associated genus *Desulfivobrio*^30, 31^ (Figure 1D) as well as a ∼4-fold increase in the *Firmicutes* to *Bacteroidetes* (F/B) ratio (Figure 1E). Interestingly, WD_CNTL_ fed mice showed negligible changes in the microbiome signature during the 6 weeks of intervention (Figure 1B-F & S1A-B), suggesting that the added lipid source had limited influence on the intestinal ecology. In contrast to this observation, we noted a pronounced shift in bacterial composition in mice fed WD_McB_. Within the first 2 weeks of treatment, the general community structure in these mice shifted towards that of their LFD-fed counterparts (Figure 1B-F, Table S3). We next asked if the observed taxonomical differences between groups related to alterations in the functional potential. SCFAs are main end products of metabolized fibers, and to a lesser extent amino acids escaping digestion in the SI, with vast impact on host physiology^32, 33^. The highest levels of SCFAs are found in the cecum and proximal colon^34^. We therefore investigated if cecal SCFA levels were different between groups. We found a consistent increase in the levels of the three major as well as three minor classes of SCFAs in the cecum of WD_McB_ fed mice compared to WD_CNTL_ fed counterparts pointing towards not just taxonomically, but also functionally, discrete microbiota profiles in the two groups of mice, supporting a beneficial health impact on dietary inclusion of McB lysates (Figure 1G-H).

### WD_McB_ feeding stimulates induction of gut-specific regulatory T cells

The intricate relationship between gut microbes and host immunity, combined with the immunoregulatory capacity of SCFAs^35^, prompted us to investigate if the observed changes mediated by WD_McB_ feeding were associated with immune alterations.

We accordingly analysed the immune cell profile of SI-LP and LI-LP in a subset of experimental mice (n = 6-10/group) using multicolour flow cytometry focusing on phenotypic characterization of group 3 innate lymphoid cells (ILC3), natural killer (NK) cells and T cells (consult Figure S2A-B for gating strategies). Numbers of ILC3s, NK cells and T cell receptor (TCR)-γδ^+^ T cells, were similar between groups (Figure S2C-E). The same was true for the numbers of TCRβ^+^ CD4^+^ T cells, as well as the proportion of T helper (T_H_)1-, T_H_17- and _n_T_regs_ cells (Figure 2A-B).

**Figure 2:**
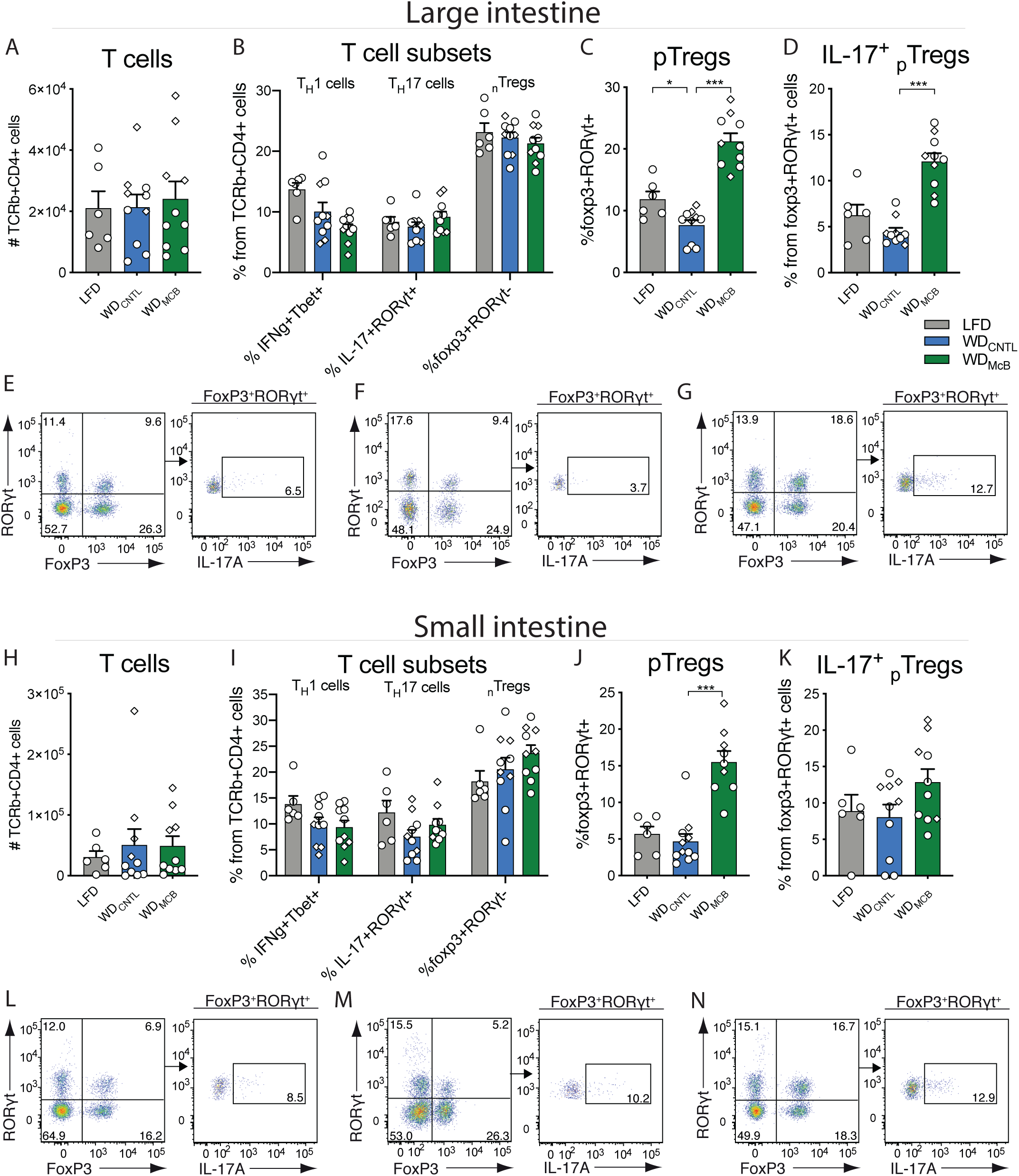
WD_McB_ feeding stimulates induction of gut-specific regulatory T cells: **A-D)** Number of indicated cells in colon from Exp1 (◆) and Exp2 (●). E-G) Representative plots of colonic TCRb^+^CD4^+^ FoxP3^+^ RORγt^+^ _p_T_regs_ (left) and IL17^+^ _p_T_regs_ (right) in LFD (E), WD_CNTL_ (F), and WD_McB_ (G) group. **H-K)** Number of indicated cells in small intestine from Exp1 (◆) and Exp2 (●). **L-N)** TCRb^+^CD4^+^ FoxP3^+^ RORγt^+^ _p_T_regs_ (left) and IL17^+^ _p_T_regs_ (right) in LFD (L), WD_CNTL_ (M), and WD_McB_ (N) group. **A-N)** n = 6 (LFD) to 10 (WD_McB_ and WD_CNTL_). **A-D, H-K)** Statistical significant differences to WD_CNTL_ group by one-way ANOVA with multiple comparisons and Dunnett post hoc is indicated by *. * = p < 0.05; ** = p < 0.01, and *** = p < 0.001.

Interestingly, the proportion of _p_T_regs_ was more than 2-fold increased in LI-LP of WD_McB_ fed mice compared to WD_CNTL_ fed counterparts (Figure 2C, p < 0.001). Notably, this regulatory T cell subset has been shown to curb intestinal inflammation^12^ and mediate immunological tolerance to the gut pathobiont *Helicobactor hepaticus*, thereby protecting against T_H_17-mediated barrier dysfunction and subsequent colitis^36^. Because RORγt is the hallmark transcription factor for T_H_17 cell differentiation and essential for their IL-17 production, we next assessed if the _p_T_regs_ induced by the different diets were also capable of expressing IL-17. Indeed, *ex vivo* stimulated LI-LP _p_T_regs_ produced substantial and diet-dependent amounts of IL-17 protein (Figure 2D-G, p < 0.001). A similar phenotype was observed in the SI-LP (Figure 2H-N), where _p_T_regs_ in WD_McB_ fed mice reached a ∼3-fold increase compared to WD_CNTL_ fed mice (Figure 2J, p < 0.001).

### WD_McB_ mitigates diet-induced obesity

The altered immune profile combined with a shift of the gut microbiota towards a state similar to that observed in lean LFD-fed mice, could potentially elicit crosstalk to glucoregulatory organs. To examine if McB lysates could reverse impaired glucose regulation, we performed OGTTs and assessed GSISs concomitant with body mass composition in obese mice fed WD_REF_ for 11 weeks and after 5 weeks of dietary intervention allowing for temporal analyses (Figure 1A). All mice were stratified into experimental groups based on their pre-intervention glucoregulatory capacity (Figure S3A-B). While the response to glucose challenge remained largely unaffected from week 11 to week 12+5, regardless of experimental diets (Figure 3A-C), both 5h fasted insulin levels and glucose stimulated insulin responses were significantly increased in mice fed WD_CNTL_ (Figure 3E, p < 0.01 & p < 0.001, respectively) in accordance with our previous report on time-dependent alterations in glucose regulation^37^. LFD- and WD_McB_-fed mice were fully protected against this detrimental trajectory (Figure 3D, F & S3C, p = 0.24 & 0.68, respectively), and WD_McB_-fed mice further exhibited modestly improved insulin sensitivity based on 5h fasted glycemia (Figure 3C, p < 0.05) and intraperitoneal insulin tolerance test (Figure 3G, p < 0.05).

**Figure 3:**
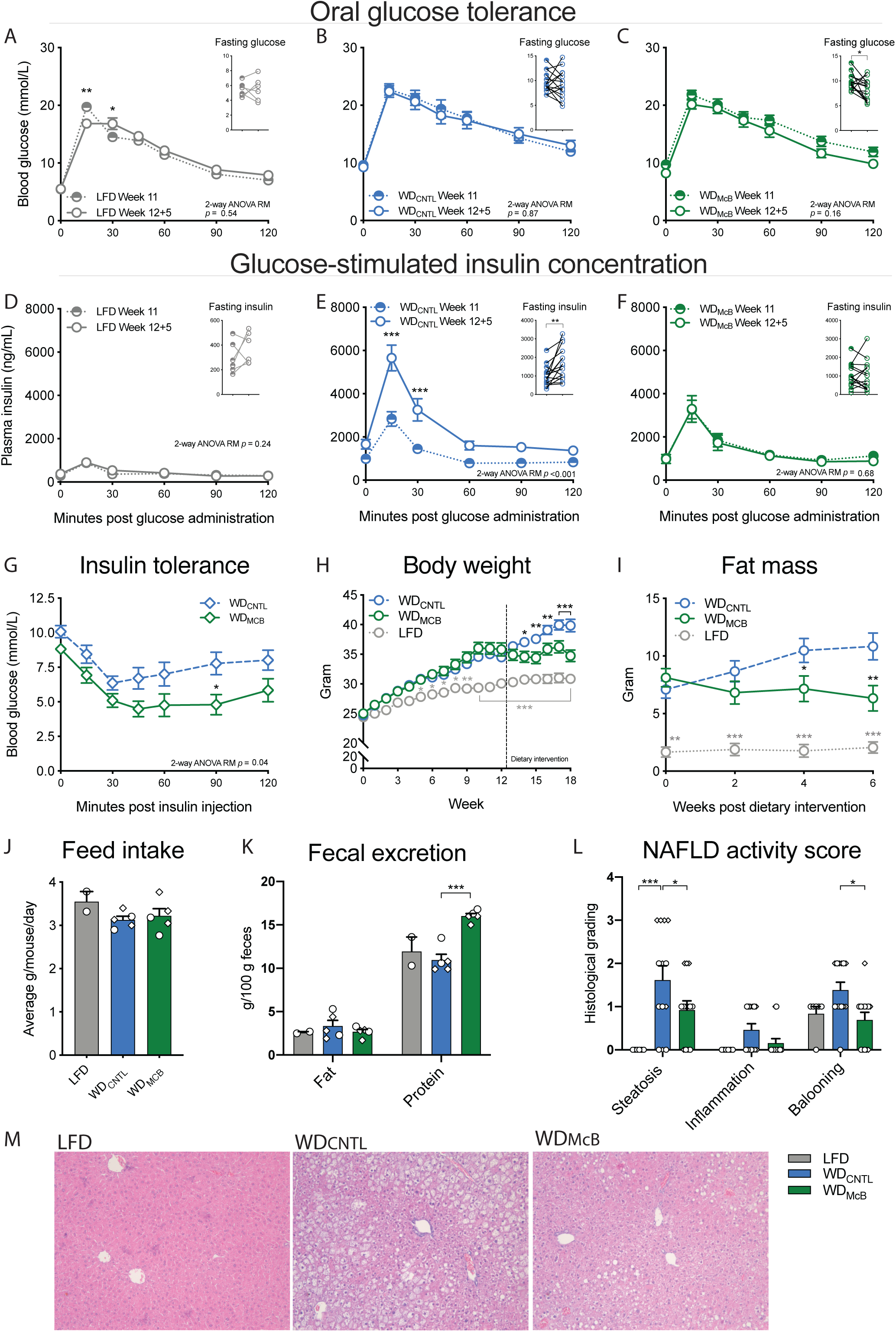
Dietary intervention with McB blunts progression of insulin resistance and fat mass accumulation: **A-C)** Oral glucose tolerance test (OGTT) and 5h fasting blood glucose prior to dietary intervention (Week 11) and 5 weeks post intervention (Week 12+5) of LFD (A), WD_CNTL_ (B), and WD_McB_ (C) groups showing mean ± SEM of two independent experiments. **D-F)** Glucose-stimulated insulin concentration during OGTT and 5h fasting insulin levels prior to dietary intervention (Week 11) and 5 weeks post intervention (Week 12+5) of LFD (D), WD_CNTL_ (E), and WD_McB_ (F) groups showing mean ± SEM of two independent experiments. **G)** Intraperitoneal insulin tolerance test (ITT) 3 weeks post dietary intervention showing mean ± SEM. **H)** Body weight development. Mice were fed either LFD or WD_REF_ the first 12 weeks followed by 6 week dietary intervention period showing mean ± SEM of two independent experiments. Dotted vertical line depicts intervention start. **I)** Fat mass in gram through the dietary intervention period (Week 12+0 to 12+6) measured by MR scan showing mean ± SEM of two independent experiments. **J)** Feed intake per cage as average grams per mouse during 48h in Exp1 (◆) and Exp2 (●). **K)** Fecal content of fat and protein in Exp1 (◆) and Exp2 (●). **L)** NAFLD activity score based on hepatic steatosis grade (0-3), inflammation (0-2) and hepatocellular ballooning (0-3) graded by blinded histological assessment of liver tissue in Exp1 (◆) and Exp2 (●). **M)** Representative H&E stain of liver tissue from indicated experimental group. **A-F)** n = 6 (LFD) to 15 (WD_McB_ and WD_CNTL_). Statistical significance within each time point is indicated by * at paired two-way ANOVA-RM with Bonferroni post hoc test. Fasting glucose and insulin levels were evaluated by paired t-tests. **G)** n = 8 (WD_McB_) to 9 (WD_CNTL_). Statistical significance within each timepoint is indicated by * during two-way ANOVA-RM with Bonferroni post hoc test. **H-I)** n = 6 (LFD) to 15 (WD_McB_ and WD_CNTL_). Statistical significance is indicated by *. LFD and WD_McB_ were compared to WD_CNTL_ by two-way ANOVA-RM, adjusted for multiple comparisons by Dunnett’s post hoc. **J-K)** Each data point resembles the average of one cage. n = 2 (LFD) to 5 (WD_McB_ and WD_CNTL_). Statistical significance is indicated by *. LFD and WD_McB_ were compared to WD_CNTL_ by two-way ANOVA-RM, adjusted for multiple comparisons by Dunnett’s post hoc. **L)** n = 6 (LFD) to 13 (WD_McB_ and WD_CNTL_). Statistical significance is indicated by *. LFD and WD_McB_ were compared to WD_CNTL_ by Kruskal-Wallis, adjusted for multiple comparisons by Dunn’s post hoc. **A-L)** * = p < 0.05; ** = p < 0.01, and *** = p < 0.001.

Overall, weight development mimicked the glucoregulatory capacity. As such, WD_McB_-fed mice exhibited stability of weight, fat mass and lean mass when changed to experimental diets, contrasting the continuous weight and fat mass development of WD_CNTL_ fed mice (Figure 3H-I & S3D). The absence of weight gain was not explained by decreased feed intake (Figure 3J), but rather appeared to be associated with enhanced fecal energy secretion (Figure 3J-K). Since obesity and impaired glucose regulation are tightly associated with NAFLD^38^, we next subjected paraffin-embedded liver sections to histological evaluation. These analyses revealed both diminished steatosis and hepatocellular ballooning in WD_McB_ fed mice compared to WD_CNTL-_fed counterparts, where especially hepatocellular ballooning was arrested in (or returned to) a state reminiscent that of lean LFD fed mice (Figure 3L-M, p < 0.05). Importantly, hepatocellular ballooning is instrumental in the development of the more severe liver disease, NASH^38^.

### WD_McB_ feeding resets the hepatic lipidome and decreases hepatic immune infiltration alleviating NAFLD

Based on the decreased NAFLD in WD_McB_-fed mice housed at T_22°C_ we designed a new experiment (study outline, Figure 4A) using a recently described^39^ method where thermoneutral housing (T_30°C_) potentiates NAFLD in WT C57BL/6J mice fed an obesogenic diet for 20-24 weeks. To more thoroughly investigate the effect of WD_McB_-feeding, we also redesigned the diets and omitted macadamia oil in the WD_CNTL_ group, as this might lead to progression of obesity (Figure 3) and related disorders. This new diet design entailed an increased fat/protein ratio in WD_McB_ compared to WD_REF_, due to phospholipids inherently present in bacterial lysates^16^ (Table S1). Despite the lower protein content in WD_McB_ compared to both other diets, protein availability was well beyond critical levels, corroborated by similar lean mass to WD_REF_-fed mice post diet intervention (Figure S4A). Still, WD_McB_-fed mice exhibited significantly improved 5h fasting insulin levels and decreased fat mass (Figure 4B-C), despite weight maintenance and significantly increased energy intake compared to both LFD and WD_REF_ fed mice (Figure S4B-D). The decreased body fat mass was accompanied by a diminished NAS, supported by both pathological evaluation of H&E stained liver sections (Figure 4D-E) and hepatocytic lipid content assessed by Oil-Red-O staining (Figure 4F-H). We additionally observed augmented adiponectin secretion (Figure 4I), pointing towards improved insulin sensitivity in the WD_McB_ group, further supported by the assessment of insulin tolerance and hepatic gene transcription activity of key metabolic enzymes (Figure 4J-K). Of interest, we observed a >10-fold downregulation of *Scd1* in the liver of WD_McB_-fed mice, the hepatic expression of which is a) regulated by the microbiota^40^ and b) instrumental in *de novo* lipogenesis at the onset of metabolic syndrome^41^.

**Figure 4:**
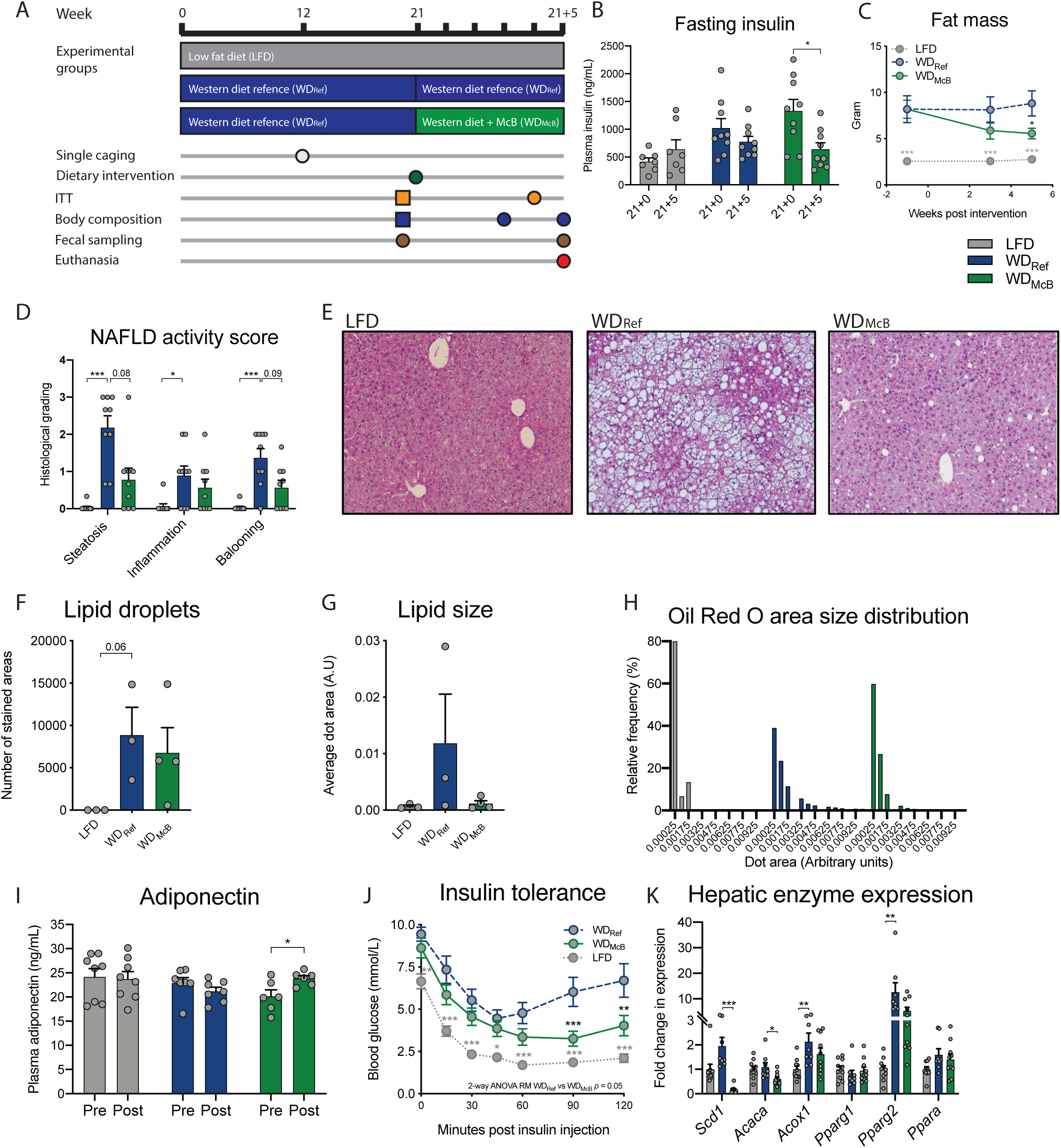
Dietary McB intervention improves diet-induced metabolic and hepatic phenotype after prolonged WD feeding: **A)** Study outline with initial Western diet (WD_REF_) feeding for 21 weeks at 30°C. Insulin tolerance test (ITT) and body composition by MR scan were assessed after 20 weeks to stratify groups (⃞) followed by dietary intervention for 5 weeks of feeding either WD_REF_ or WD including McB lysate (WD_McB_). A reference group fed low fat diet (LFD) was included throughout the experiment (week 0 to week 21+5). Body composition, ITT, and fresh fecal sampling was carried out at the indicated intervals. Mice were co-housed the first 12 weeks and then separated to individual cages. **B)** 5h fasting plasma insulin levels before (Week 20) and after (Week 21+5) dietary intervention. **C)** Fat mass in grams measured by MR scan at indicated time points from dietary intervention. **D)** NAFLD activity score separated in hepatic steatosis grade (0-3), inflammation (0-2) and hepatocellular ballooning (0-3) graded by blinded histological assessment of H&E stained liver tissue. **E)** Representative H&E stain of liver tissue from indicated experimental group. **F)** Number of lipid droplets in each of 3-4 liver samples per experimental group quantified by Oil Red O staining. **G)** Average size of lipid droplets in each of 3-4 liver samples per experimental group quantified by Oil Red O staining. **H)** Distribution of lipid droplet size in % of all lipid droplets within each experimental group. **I)** Plasma adiponectin concentration in 5h fasted mice before (Week 20) and after (Week 21+5) dietary intervention. **J)** ITT after four weeks of dietary intervention (Week 21+4). **K)** Expression of key metabolic enzymes in liver tissue after five weeks of dietary intervention (Week 21+5) by RT-qPCR. **B+I)** n = 6-9 per group. Statistical significance is indicated by * on paired t-test. **D+K)** n = 8-10 per group. Statistical significance compared to WD_REF_ by Kruskal-Wallis test, adjusted for multiple comparisons by Dunn’s post hoc. **F-H)** n = 3 randomly selected samples per group. Statistical significance compared to WD_REF_ by Kruskal-Wallis test, adjusted for multiple comparisons by Dunn’s post hoc. **C+J)** n = 9-10 per group. Statistical significance compared to WD_REF_ by two-way ANOVA-RM, adjusted for multiple comparisons by Dunnett post hoc. * = p < 0.05; ** = p < 0.01, and *** = p < 0.001.

We next assessed the hepatic lipidome by mass spectrometry (MS/MS) to elucidate if diminished NAFLD was associated with an altered lipid profile. Through comparison of WD_McB_ and WD_REF_ we identified 57 and 279 differentially regulated peaks in negative and positive ionization mode, respectively (Figure 5A-B, S5A-B, all FDR < 0.05). Of these, most classified lipids were changed with WD_McB_ in the direction of LFD fed mice (Figure 5C-D). Notably, 57% of upregulated species were odd-chain fatty acids, whereas 80% of down-regulated species represented lipids with even carbon numbers (Figure 5C-D & Table S4), hence supporting previous reports where odd-rather than even-chain fatty acids are inversely associated with human insulin resistance^9^ and type 2 diabetes^42^.

**Figure 5:**
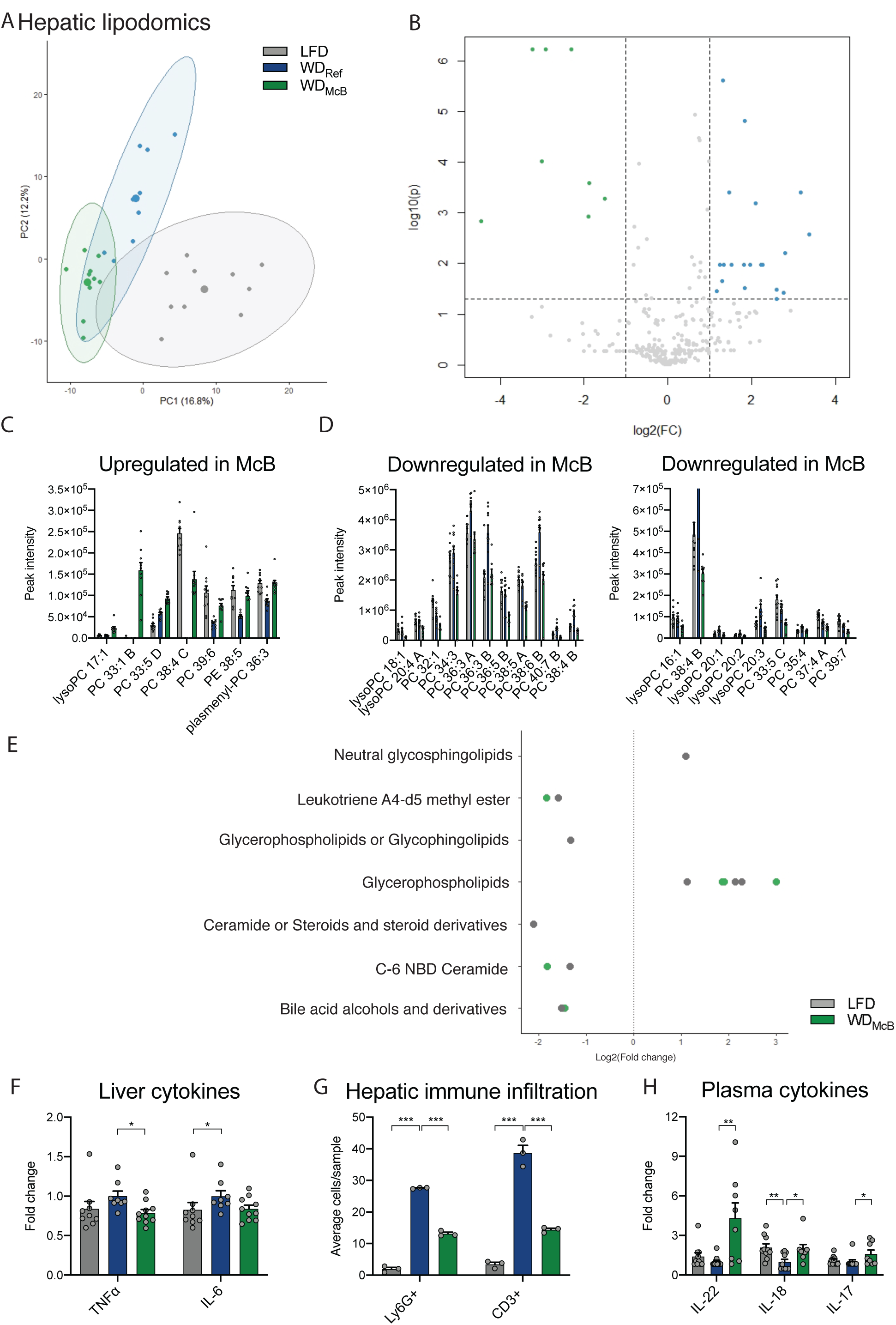
WD_McB_ feeding resets the hepatic lipidome and decreases hepatic immune infiltration alleviating NAFLD: **A)** Principle coordinate analysis (PCoA) of hepatic lipid species identified in negative ionization mode. **B)** As in A but depicted in a volcano plot with significantly regulated lipid species (FDR < 0.05) presented on a log2 scale in either green (WD_McB_ > WD_REF_) or blue (WD_McB_ < WD_REF_). **C-D)** Identified lipid species differentially expressed (FDR < 0.01) between WD_McB_ and WD_REF_ as indicated. Fold-change and adjusted p values of individual lipid species are indicated in Table S4. **E)** Lipd pathways identified by fold-change analysis seperating LFD (grey) and WD_McB_ (green) groups from WD_REF_ in negative ionization mode. **F)** Hepatic cytokine levels of TNF-α and IL-6 as indicated. **G)** Ly6G^+^ and CD3^+^ immune cell revealed by immunohistochemistry. **H)** Cytokine levels in plasma of IL-22, IL-18, and IL-17. **F+H)** n = 8-10. Statistical significance compared to WD_REF_ group by Kruskal-Wallis test, adjusted for multiple comparisons by Dunn’s post hoc. **G)** n = 3 per group (randomly selected). Statistical significance compared to WD_REF_ by one-way ANOVA, adjusted for multiple comparisons by Dunnett post hoc. * = p < 0.05; ** = p < 0.01, and *** = p < 0.001.

To estimate the functional consequences of an altered lipid profile, we used the Lipidmaps database to identify affected pathways and plotted the observed changes on a log_2_ scale comparing both LFD and WD_McB_ to WD_REF_. The majority of affected pathways were similarly regulated in both direction and magnitude in LFD and WD_McB_ mice compared to WD_REF_ mice (Figure 5E & S5C). Of note, bile acids and ceramides, both of which were significantly downregulated in WD_McB_ fed mice compared to WD_REF_ fed counterparts, have been shown to mediate steatohepatitis by upregulation of IL-6 and TNF-α, respectively^43, 44^. We therefore measured these hepatic cytokines and observed similarly reduced levels in both LFD and WD_McB_ compared to WD_REF_ (Figure 5F).

A key feature of diet-induced liver pathologies, including NAFLD, is recruitment of newly activated immune cells capable of eliciting a proinflammatory immune response. This process is generally hampered in mice housed at mild thermal stress, which therefore fail to phenocopy human pathophysiology. However, thermoneutral housing recapitulates some human disease traits^45^, which combined with HFD-feeding accentuates intrahepatic infiltration of proinflammatory Ly6^high^ monocytes^46^. These monocytes interact with tissue resident T cells and play a central role in the pathogenesis of liver injury, hence representing an attractive therapeutic target to mitigate NAFLD development and to curtail associated pathologies^44^.

We therefore subjected liver tissues from representative mice to immunological evaluation by immunohistochemistry and observed a marked decrease in both CD3^+^ T cells and Ly6^high^ monocytes in WD_McB_-fed mice compared to their WD_REF_-fed counterparts (Figure 5G). Diminished hepatic immune infiltration was mirrored by increased levels of circulating IL-22, IL-18, and IL-17 in WD_McB_-fed mice compared to their WD_REF_-fed counterparts (Figure 5H, p < 0.01, < 0.05 & < 0.05, respectively).

### WD_McB_ feeding reverses prolonged gut microbial dysbiosis and markedly improves colonic mucus production

The improved hepatic phenotype prompted us to further investigate potential traits in the gut-liver axis. We initially assessed if the observed cytokine responses associated with improved gut health in mice housed at T_30°C_, where inflammation is expected to be increased^45^. Indeed, WD_McB_-fed mice were resistant to WD-induced colonic shortening closely associated with colonic inflammation (Figure 6A).

**Figure 6:**
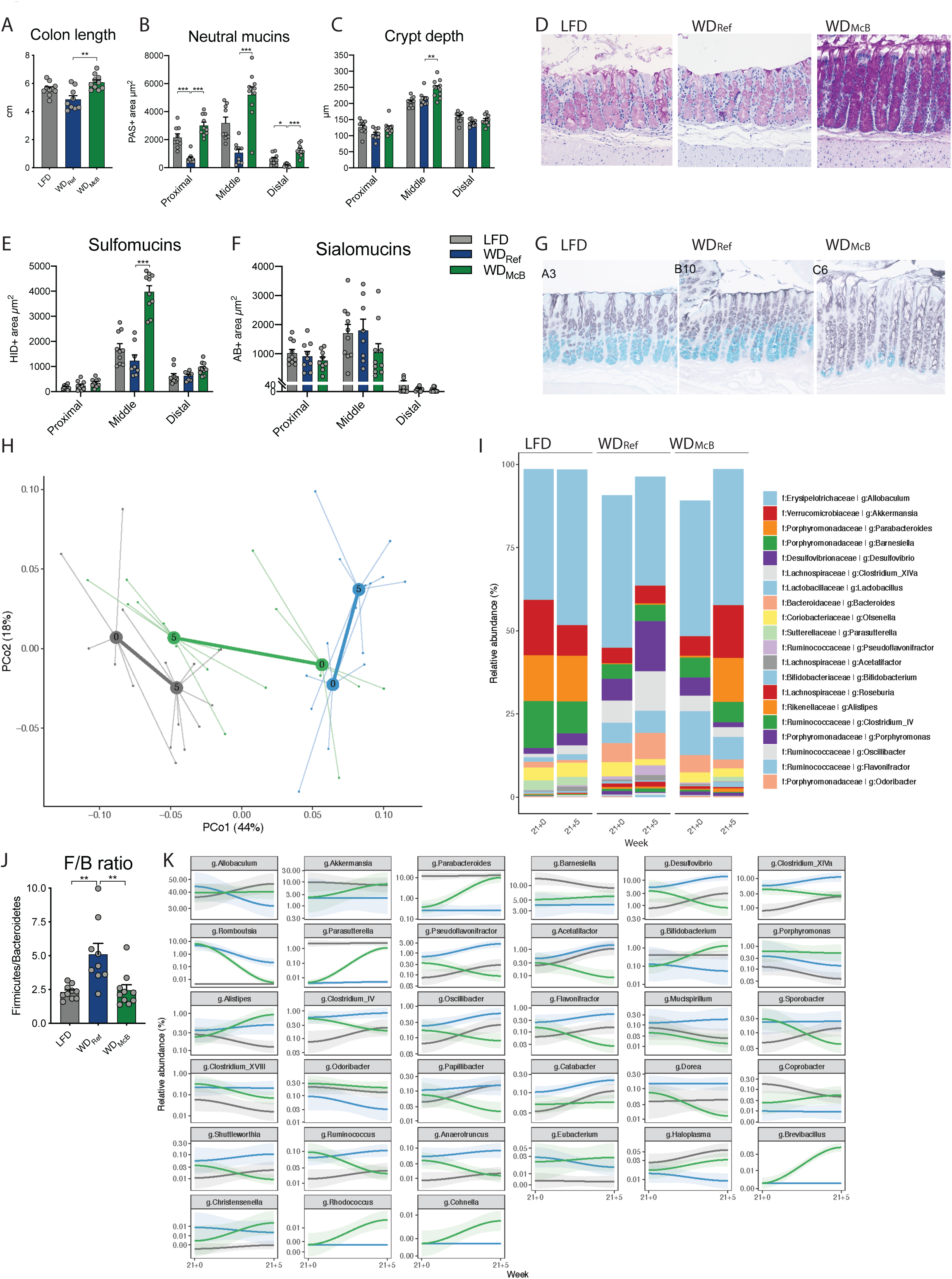
WD_McB_ treatment improves colonic mucus production and reverses obesity-induced gut microbiota changes to resemble the composition found in lean LFD-fed mice: **A)** Colon length in centimeters. **B)** Area of neutral mucins determined by Periodic Acid Schiffs (PAS) staining in three descrete locations, i.e. proximal, middle, and distal colon segments. Three longitudinally sectioned crypts were measured in each sample (one data point) at each location. **C)** Colon crypt depth (CD) in proximal, middle, and distal colon segments, measured in histological sections. Measurements are given in µm, and results are given as means of six crypt measurements per location per individual mice (3 longitudinally PAS stained and HID-AB stained crypts, respectively, per data point). **D)** PAS staining for mucus in colon middle segments, representative histological sections from the three experimental groups as indicated. **E)** Area of sulfomucins in proximal, middle, and distal colon segments by HID staining. Three longitudinally sectioned crypts were measured in each sample and at each location. **F)** Area of sialomucins in proximal, middle, and distal colon segments by AB staining. Three longitudinally sectioned crypts were measured in each sample and at each location. **G)** Representative HID-AB staining of middle segment of indicated experimental group. **H)** PCoA of fecal microbiota composition of indicated group before (Week 21+0) and after (21+5) dietary intervention with group mean indicated as centroids. Microbiota composition was significantly different between the WD_REF_ and WD_McB_ groups at the end of the experiment week 21+5 (PERMANOVA p = 0.001) **I)** Taxasummary of most abundant bacterial genera showing mean relative abundance of indicated genera in each group at indicated time point. **J)** Firmicutes/Bacteroidetes ratio of fecal samples of individual mice at termination (Week 21+5). **K)** Deseq analysis of fecal bacterial genera abundances significantly regulated by McB intervention (p.adj. < 0.05). Relative abundance in % in each group and variation are shown for each regulated genera at the sampled time points. Fold-change and adjusted p values of individual genera are indicated in Table S5. **A-K**) n = 8-10. **A-C, E-J)** Statistical significance compared to the WD_REF_ group by one-way ANOVA or Kruskal-Wallis test (depending on Gaussian distribution) with multiple comparisons by Dunnett or Dunn’s post hoc, respectively. * = p < 0.05; ** = p < 0.01, and *** = p < 0.001.

To gain additional insights to the immunomodulating properties of McB lysates, we focused on mucus production and -function. Because mucin production is a constitutive process where both secretion and adherence are constantly ongoing and rapidly adjust to environmental changes, immunohistochemical labelling for various MUC epitopes may not fully recapitulate the physical properties of the mucus. We therefore applied specialized mucin histochemistry staining allowing us to differentiate between neutral and acidic mucins according to the net charge of each molecule. Acidic mucins were further separated into sulfomucins and sialomucins. Sections revealed that neutral mucins in crypt-residing goblet cells were consistently down regulated in WD_REF_-fed mice compared to LFD-fed counterparts in three well defined segments of the colon; i.e. proximal, middle and distal area (Figure 6B, D). WD_McB_-feeding not only reversed this pattern in all three segments but even enhanced the production of neutral mucins exceeding the levels found in LFD-fed counterparts. This is a remarkable finding considering the continuous intake of a westernized diet high in fat and sucrose, known to hamper goblet cell function. We next evaluated crypt depth (CD) in stained sections. While the CD of the proximal and distal part of the colon were largely unaffected by diet, we observed increased CD in the middle segment of WD_McB_-fed mice (Figure 6C). WD_McB_-feeding further enhanced the glycosylation pattern, particularly in the middle segment of colon, where this group exhibited a 3-fold increase in sulfomucins balanced by a similar (∼2-fold) decrease in sialomucins compared to WD_REF_-fed mice (Figure 6E-G). No differences were observed between WD_REF_ and LFD-fed mice, indicating that the reciprocal regulation of mucin glycosylation status was specific to WD_McB_-feeding.

With focus on the dynamic interactions between gut immunity, mucin glycosylation and commensal microbes^47^, we next assessed the gut microbiota composition in temporally separated samples. This was done to determine if the changes towards a gut microbiota resembling that of lean LFD-fed mice was recapitulated in this intensified setup. In contrast to the first set of experiments, where we used cohoused mice shown to exhibit resilient microbiota profiles, we now employed single-housed mice to explore if the WD_McB_-mediated community structures were persistent enough to induce consistent changes in the more dynamic communities of single-housed mice. Similar to our first experiments at T_22°C_, we observed a normalization of the gut microbiome of WD_McB_-fed mice, despite prolonged WD-feeding prior to intervention (Figure 6H-K). WD_McB_-induced changes were surprisingly consistent with the first set of experiments, including a significantly lower F/B ratio (Figure 6J and S5D), and a substantial reduction of *Desulfovibrio* abundance, countered by a similar bloom of the *Parasutterella* and *Parabacteroides* genera (Figure 6K, Table S5). Of note, the age-related increases in *Desulfovibrio* abundances recently reported^30^ was confirmed in this study where the general magnitude in both WD_REF_- and LFD-fed mice increased 2-3-fold over the 5 week intervention (Figure 6K). WD_McB_-feeding fully prevented this trajectory and paired analyses even revealed a diminished relative abundance of *Desulfovibrio* in these mice.

## Discussion

In this report, we explored the relationship between dietary nutrients and host-microbe interactions with a focus on immunometabolic response rates in the context of high fat, high sucrose, WD-feeding. We found whole-cell lysates from the non-commensal methanotrophic bacterium, McB, capable of reversing hallmark signatures of WD-feeding despite continued intake of an obesogenic diet. The signatures included dimished fat mass, improved intestinal immunity and glucoregulatory capacity accompanied by gut microbial community structures resembling that of lean LFD-fed counterparts.

The aberrant microbiota composition associated with obesity was recently shown to reflect HFD intake rather than obesity *per se*^48^. It is therefore worth noting that we were able to normalize the dysregulated gut microbiome in mice remaining on a westernized, high fat, high sucrose diet; especially considering that it remains a clinical challenge to change fatty dietary habits in lifestyle-related obese subjects. This remarkable change was reproduced in two diverse sub-strains of mice originating from different vendors, corroborating a robust phenotype not notably affected by baseline microbiota composition. To this end, a previous report on the resilient microbiome, argued for prolonged normalization^49^. This study found that mice transferred to a low fat, fiber-rich chow diet after 12 weeks of HFD feeding, shifted their microbiota towards age-matched chow-fed control mice within 4 weeks, but only fully converged 10 weeks post diet change. This is particularly interesting as dietary fibers are known to be the most potent dietary regulator of the gut microbiota^50^, by far exceeding that of dietary fat^51^. Still, in our hands, WD_McB_-feeding was able to reverse the obese microbiome at a higher pace than chow diet was in the previous report^49^, despite similar fiber content in the two WDs. We further showed that the reversal of the microbiota traits were reproducible at different temperatures, altered reference diets, changed experimental duration, and in both co-housed and single-housed mice; all of which are prominent modulators of gut microbiota community structures. While many genera were similarly affected between experiments, others were either exclusively regulated at T_30°C_ (e.g., *Bifidobacterium*) or less pronounced affected at T_30°C_ (e.g., *Barnesiella*), suggesting dispensability for these specific microbes in the metabolic disease traits observed in this model. Similar observations were made for *Akkermansia*, which was ∼3-fold upregulated at T_30°C_, but discordantly affected (log2 FC bouncing from +9.3 to -10.4) between the two replication studies at T_22°C_ with pronounced intragroup variation.

Contrasting these discrepancies, there was a consistent downregulation of *Desulfovibrio* accompanied by >10-fold upregulation of *Parabacteroides* and *Parasutterella* suggesting that WD_McB_ feeding nourishes a favorable microbiota, albeit such speculations have yet to be mechanistically described. Still, *Desulfovibrio* was most recently shown to flourish in aged, immunocompromised, obese mice with impaired glucose regulation^30^. Both T cell loss and *Desulfovibrio* administration *per se*, precipitated obesity-induced insulin resistance^30^. This bacterium has moreover been shown to thrive in desulfonated colonic mucosa, associated with human^52^ and mouse^53^ colitis. Combined, these observations lend credence to the hypothesis of a mechanistic link between WD_McB_-mediated reductions of *Desulfovibrio* and the observed improvements in mucin chemotype and metabolic response rates.

Future studies are warranted to elucidate if the gut microbiota changes observed in WD_McB_-fed mice were a result of the unique nutrient source, or if it was facilitated by WD_McB_-induced _p_T_regs_. On this note, mucin-producing goblet cells are capable of delivering luminal antigens to LP-residing dendritic cells (DC)^54^ instrumental for T cell polarization^55^. As McB potently induce DC maturation markers *in vitro* – even exceeding the effects of the well-described probiotic strain, *Escherichia coli* Nissle 1917^56^ – we predict a direct link between McB intake and the corresponding immune profile. In addition to the direct link proposed here, McB intake might also indirectly promote T_reg_ polarization through augmented SCFAs^57^. These important metabolites also sustain mucus production and facilitate tissue crosstalk in the gut-liver axis^2^. In keeping with this notion, we observed decreased hepatic bile acids and TNF-α levels combined with a pronounced reduction of intrahepatic CD3^+^ and Ly-6C^high^ cells. Importantly, while Ly-6C^high^ cells represent proinflammatory, fibrogenic macrophages^58^, their *in situ* differentiation to Ly-6C^low^ cells^59^ facilitate tissue repair and improved NASH prognosis^60^. Mentioned targets are all proposed as relevant strategies to curb NAFLD and the more terminal liver disease, NASH^44^.

The substantial, consistent, and global changes in the immunometabolic profile found in this study point towards clinical potential if further developed. Notably, only a single study has previously succeeded in inducing _p_T_regs_, and this was limited to LI-LP^11^. Apart from inducing this cell subset throughout the gastrointestinal tract, we further show that these cells exhibited enhanced secretory capacity of IL-17 in WD_McB_-fed mice compared to their WD_CNTL_ fed counterparts. To this end, emerging evidence indicates that the non-regulatory counterparts to _p_T_regs_, i.e. T_H_17 cells, are purged from SI-LP in DIO mice^14^ and that reintroducing *ex vivo* differentiated gut tropic T_H_17 cells curtail obesity development, hence improving insulin sensitivity^15^.

That IL-17 in this intervention is produced by T_regs_ and not T_H_17 cells might further improve the safety profile of a putative medication, as the physiological impact and properties of T_H_17 cells are context-dependent. As such, T_H_17 cells may potentiate inflammatory bowel disease (IBD) in individuals with compromised barrier function^61^, whereas _p_T_regs_ have shown enhanced suppressive capacity against the same disease^12^. This is a pertinent feature given the relatively high proportion of subjects with metabolic complications co-suffering from IBD^62^. While our study was not designed to assess IBD susceptibility, it is well described that purified diets compromise barrier function^50, 63, 64^ and thus represent clinical IBD-features (disease-cause) preceding inflammation (symptom)^65^. It is therefore worth noting that in addition to the immunomodulating phenotype, WD_McB_-feeding augmented neutral mucus production in all segments of the colon while enhancing IBD-protective sulfomucins specifically in the middle segment. In this segment, we also detected increased CD in McB fed mice, collectively pointing towards increased barrier function.

In summary, this study demonstrates a consistent activation of Foxp3^+^RORγt^+^IL-17^+^ triple positive pT_regs_ throughout the gastrointestinal tract and a reversion of gut dysbiosis in response to WD_McB_-feeding, even in the context of high dietary fat and sucrose intake. The prospects of using bacterial lysates as an alternative to traditional live probiotics merit further investigation, just as future studies are warranted to assess to what extent our results can be translated to humans. Future studies are also urgently needed to identify the biological effector molecule(s) comprised in this bacterial lysate, allowing for subsequent development of a versatile medical product(s) rebalancing intestinal immunity while targeting gut-related dysbiosis and metabolic abnormalities.

---

**Supplementary figure S1: A)** Taxasummary of most abundant bacterial genera showing mean relative abundance in % of indicated family and genera in each group at indicated time point. B) Deseq analysis of fecal bacterial genera abundances significantly regulated by McB intervention compared to the WD_CNTL_ (p.adj. < 0.05). Relative abundance in % in each group and variation are shown for each regulated genus at the sampled time points. Fold-change and adjusted p values of individual are indicated in Table S3.

**Supplementary figure S2: A)** Gating strategy for group 3 innate lymphoid cells (ILC3). Representative flow cytometry plots of *ex vivo* stimulated small intestine (SI) lamina propria (LP) cells. ILC3s were gated as single, live cells, lineage1^-^lineage2^-^CD90.2^+^CD127int/^+^RORγδ^+^ cells. Representative data of RORγδ+ ILC3s after stimulation with or without (w/o) IL-23 are shown. Lineage1: CD11c, CD19, CD45R/B220, Ly-6G/Ly-6C (Gr-1), Ter119, NK1.1; Lineage2: CD3ε, CD8α, TCRβ, TCRγδ. **B)** Gating strategy for NK cells and T cells. Representative flow cytometry plots of *ex vivo* stimulated colon lamina propria cells. NK cells were gated as single, live cells, CD8α^-^ CD45^+^NK1.1^+^; TCRγδ^+^ T cells were gated as single, live cells, CD8α^-^CD45^+^TCRγδ^+^CD4^-^; CD4^+^ T_H_1 cells were gated as single, live cells, CD8α^-^CD45^+^TCRβ^+^CD4^+^T-bet+IFNγ^+^; CD4^+^T_H_17 cells were gated as single, live cells, CD8α^-^CD45^+^TCRβ^+^CD4^+^RORγδ^+^IL-17A^+^. Natural regulatory T cells (_n_T_regs_) were gated as single, live cells, CD8α^-^CD45^+^TCRβ^+^CD4^+^FoxP3^+^RORγδ^-^ and peripheral-induced regulatory T cells (_p_T_regs_) were gated as single, live cells, CD8α^-^CD45^+^TCRβ^+^CD4^+^FoxP3^+^RORγδ^+^. Representative data of TCRβ^+^CD4^+^ T cells after stimulation with or without (w/o) PMA and Ionomycin (PMA/Iono) are shown. **C)** ILC3 cell number in large intestine (LI)-LP and SI-LP. **D)** NK1.1 cell number in LI-LP and SI-LP. **E)** TCRγδ^+^CD4^-^ cell number in LI-LP and SI-LP. **C-E)** Individual values from Exp1 (◆) and Exp2 (●) are shown and statistical significant differences to WD_CNTL_ by one-way ANOVA, adjusted for multiple comparisons by Dunnett’s post hoc. * = p < 0.05; ** = p < 0.01, and *** = p < 0.001.

**Supplementary figure S3: A-B)** Blood glucose and plasma insulin values, respectively, during oral glucose tolerance test (OGTT) before dietary intervention (week 11). Mean ± SEM is shown for two, independent experiments. **C)** Plasma insulin values from OGTT week 12+5 depicted as mean ± SEM of two, independent experiments. **D)** Lean mass in grams during the dietary intervention period as mean ± SEM of two, independent experiments. A-D) Statistical significance compared to WD_CNTL_ by two-way ANOVA-RM, adjusted for multiple comparisons by Dunnett post hoc. * = p < 0.05; ** = p < 0.01, and *** = p < 0.001.

**Supplementary figure S4: A)** Lean mass in grams during the dietary intervention period as mean ± SEM. Statistical significance compared to WD_REF_ by two-way ANOVA-RM, adjusted for multiple comparisons by Dunnett post hoc. **B)** Body weight development; mice were fed either LFD or WD_REF_ for the first 21 weeks, as indicated. WD_REF_-fed mice were subsequently stratified into experimental groups, fed indicated diets for additional 5 weeks. **C)** Final body weight week 21+5. **D)** Cumulative energy intake in kcal per mouse during dietary intervention (week 21+0 to 21+5). **C-D)** Statistically significant differences to WD_CNTL_ by one-way ANOVA, adjusted for multiple comparisons by Dunnett’s post hoc. * = p < 0.05; ** = p < 0.01, and *** = p < 0.001.

**Supplementary figure S5: A)** Principle coordinate analysis (PCoA) of hepatic lipid species identified in positive ionization mode. **B)** As in A but depicted in a volcano plot with significantly regulated lipid species (FDR < 0.05) presented on a log2 scale in either green (WD_McB_ > WD_Ref_) or blue (WD_McB_ < WD_Ref_). **C)** Lipid pathways identified by fold-change analysis that segregated LFD-(grey) and WD_McB_ (green) groups from WD_Ref_ in positive ionization mode. **D)** Firmicutes/Bacteroidetes ratio of fecal bacteria sampled prior to dietary intervention in week 21+0. Statistically significant differences to WD_Ref_ by one-way ANOVA, adjusted for multiple comparisons by Dunnett’s post hoc. * = p < 0.05; ** = p < 0.01, and *** = p < 0.001.

**Table S1:**
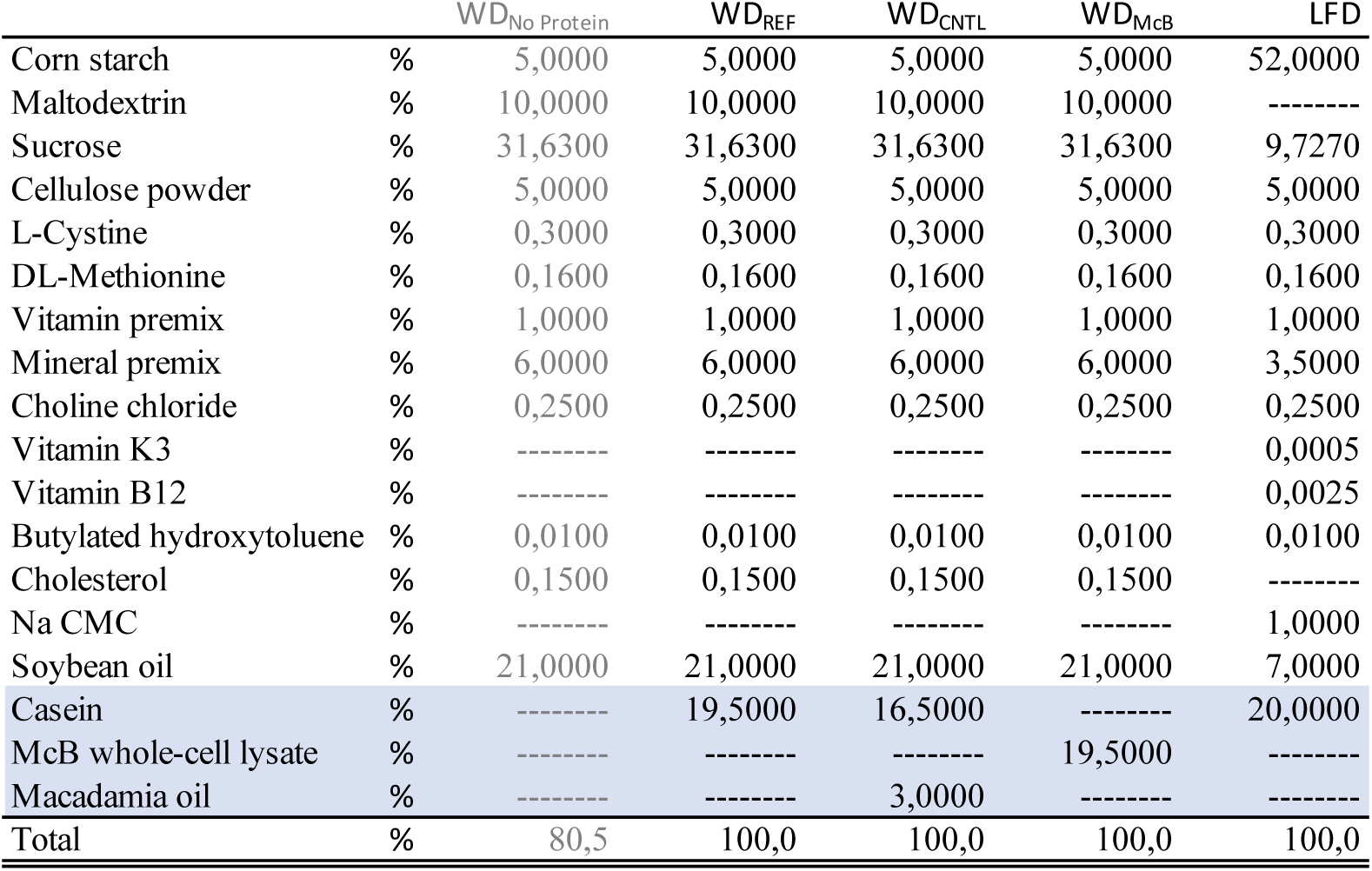
Experimental diet compositions. Composition of experimental diets used in the studies as % weight. Experimental diets were based on lot-matched WDNo Protein, and subsequently added casein, macadamia oil and McB whole-cell lysate as indicated.

**Table S2:**
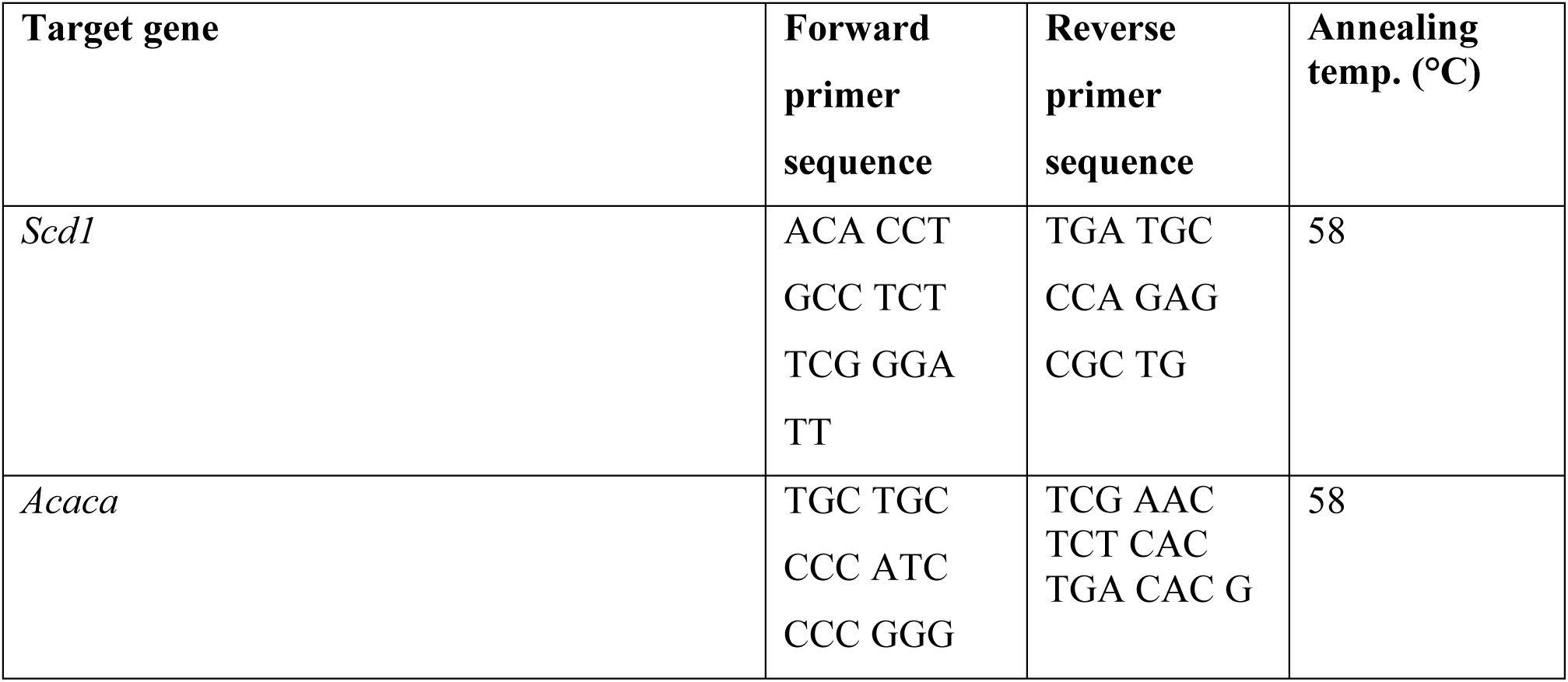
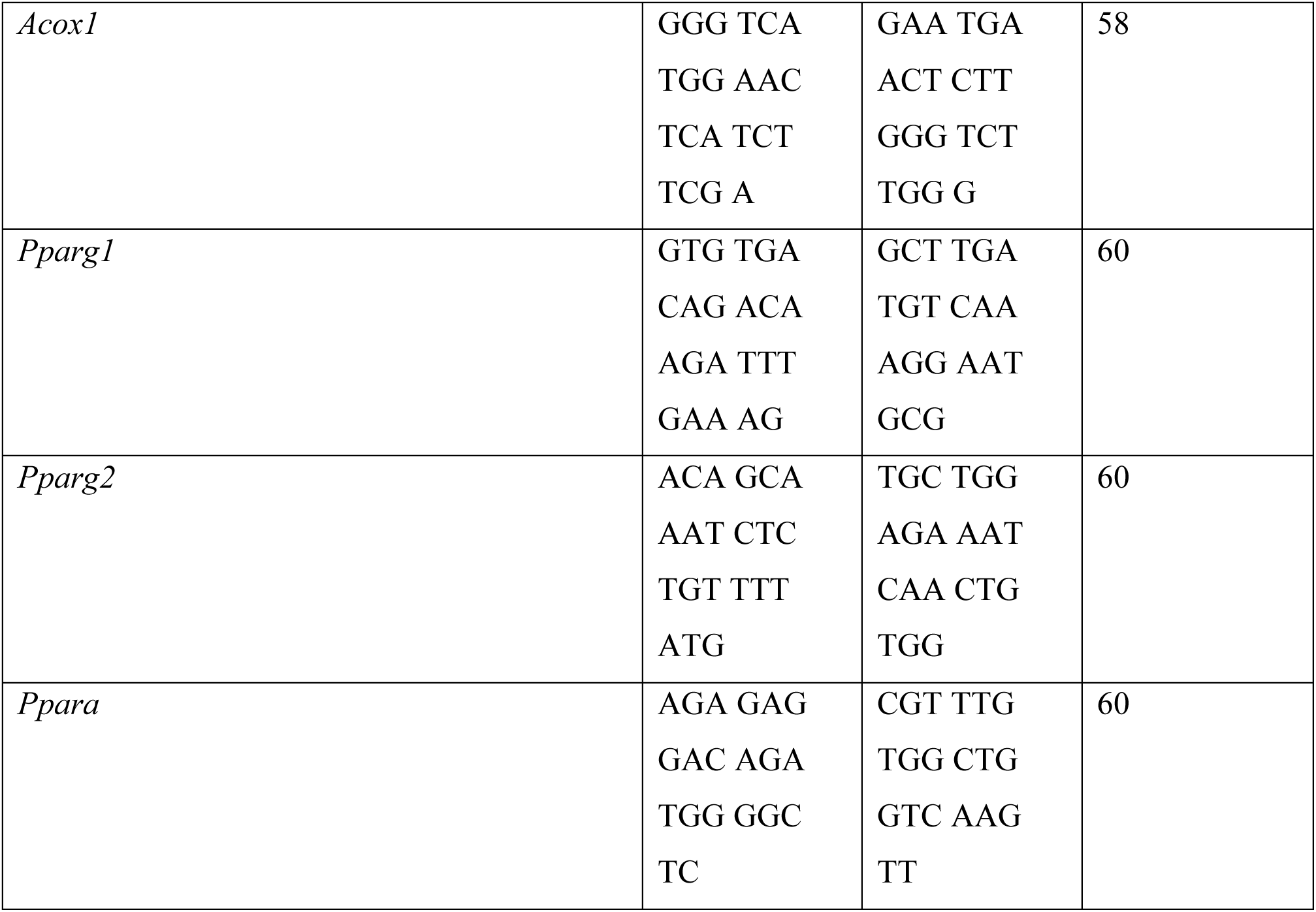
Primer sequences and annealing temperature for qPCR.

**Table S3:** Deseq analysis from 12+6 week experiments. Results from deseq analysis of bacterial genera of the two experiments presented in Figure 1 & S1 featuring genus name, Fold change and Log2 fold change, comparing WD_McB_ to WD_CNTL_. Unadjusted p-values as well as p-values adjusted by Benjamini-Hochberg method are reported and statistically significantly affected genera (Adj. p < 0.05) are marked in bold.

| Related to Figure 1 |  |  |  |  |
| --- | --- | --- | --- | --- |
| Genus | Fold change | Log2FC | p value | Adj p value |
| **g:Chelatococcus** | **1573,07** | **10,62** | **4,43E-25** | **2,88E-23** |
| **g:Barnesiella** | **27,46** | **4,78** | **3,07E-20** | **6,65E-19** |
| **g:Parabacteroides** | **23,89** | **4,58** | **2,22E-20** | **6,65E-19** |
| **g:Vampirovibrio** | **1401,99** | **10,45** | **3,08E-18** | **5,01E-17** |
| **g:Anaerofilum** | **408,29** | **8,67** | **6,55E-18** | **8,52E-17** |
| **g:Parasutterella** | **109,80** | **6,78** | **8,74E-16** | **9,47E-15** |
| **g:Microvirga** | **268,76** | **8,07** | **1,35E-14** | **1,25E-13** |
| **g:Brevibacillus** | **88,56** | **6,47** | **9,74E-09** | **7,36E-08** |
| **g:Pseudoxanthomonas** | **84,23** | **6,40** | **1,02E-08** | **7,36E-08** |
| **g:Clostridium_IV** | **0,24** | **-2,06** | **5,16E-08** | **3,35E-07** |
| **g:Isoptricola** | **51,65** | **5,69** | **5,17E-07** | **3,05E-06** |
| **g:Eubacterium** | **7,88** | **2,98** | **3,56E-06** | **1,93E-05** |
| **g:Akkermansia** | **632,32** | **9,30** | **7,69E-06** | **3,84E-05** |
| **g:Clostridium_XIVa** | **2,64** | **1,40** | **2,23E-05** | **0,0001** |
| **g:Staphylococcus** | **7,28** | **2,86** | **0,0002** | **0,001** |
| **g:Papillibacter** | **0,25** | **-2,01** | **0,0002** | **0,001** |
| **g:Acetatifactor** | **0,44** | **-1,19** | **0,001** | **0,002** |
| **g:Lachnospiracea_incertae_sedis** | **0,16** | **-2,61** | **0,001** | **0,002** |
| **g:Odoribacter** | **8,16** | **3,03** | **0,001** | **0,002** |
| **g:Bacteroides** | **2,81** | **1,49** | **0,002** | **0,01** |
| **g:Alistipes** | **2,83** | **1,50** | **0,003** | **0,01** |
| **g:Porphyromonas** | **4,77** | **2,25** | **0,004** | **0,01** |
| **g:Haloplasma** | **5,41** | **2,44** | **0,01** | **0,02** |
| **g:Desulfovibrio** | **0,44** | **-1,17** | **0,01** | **0,03** |
| **g:Blautia** | **0,24** | **-2,05** | **0,01** | **0,03** |
| **g:Escherichia/Shigella** | **0,14** | **-2,88** | **0,01** | **0,04** |
| **g:Murimonas** | **0,34** | **-1,57** | **0,02** | **0,04** |
| **g:Anaerovorax** | **3,76** | **1,91** | **0,02** | **0,04** |
| **g:Shuttleworthia** | **0,27** | **-1,87** | **0,02** | **0,05** |
| g:Alkaliphilus | 0,15 | -2,77 | 0,03 | 0,06 |
| g:Dorea | 0,41 | -1,28 | 0,03 | 0,07 |
| g:Allobaculum | 2,17 | 1,12 | 0,04 | 0,07 |
| g:Christensenella | 8,11 | 3,02 | 0,04 | 0,07 |
| g:Streptococcus | 0,34 | -1,57 | 0,03 | 0,07 |
| g:Clostridium XVIII | 0,39 | -1,36 | 0,04 | 0,07 |
| g:Clostridium III | 2,16 | 1,11 | 0,04 | 0,07 |
| g:Ruminococcus | 0,42 | -1,24 | 0,04 | 0,07 |
| g:Pseudoflavonifractor | 0,55 | -0,86 | 0,05 | 0,08 |
| g:Saccharofermentans | 2,46 | 1,30 | 0,05 | 0,09 |
| g:Lutispora | 4,23 | 2,08 | 0,06 | 0,10 |
| g:Butyrivibrio | 1,75 | 0,81 | 0,08 | 0,12 |
| g:Aestuariispira | 3,27 | 1,71 | 0,09 | 0,15 |
| g:Hydrogenoanaerobacterium | 3,16 | 1,66 | 0,11 | 0,16 |
| g:Anaerotruncus | 0,55 | -0,87 | 0,12 | 0,18 |
| g:Enterorhabdus | 0,46 | -1,13 | 0,13 | 0,19 |
| g:Flavonifractor | 0,59 | -0,76 | 0,14 | 0,19 |
| g:Roseburia | 0,54 | -0,88 | 0,14 | 0,20 |
| g:Sporobacter | 1,95 | 0,96 | 0,15 | 0,21 |
| g:Coproccoccus | 0,32 | -1,62 | 0,17 | 0,22 |
| g:Romboutsia | 0,26 | -1,95 | 0,26 | 0,34 |
| g:Intestinimonas | 0,59 | -0,77 | 0,29 | 0,36 |
| g:Butyricicoccus | 0,34 | -1,55 | 0,30 | 0,38 |
| g:Marvinbryantia | 1,69 | 0,75 | 0,35 | 0,44 |
| g:Clostridium XIVb | 0,78 | -0,35 | 0,38 | 0,46 |
| g:Clostridium sensu stricto | 0,51 | -0,97 | 0,50 | 0,59 |
| g:Gracilibacter | 1,44 | 0,53 | 0,52 | 0,60 |
| g:Catabacter | 1,22 | 0,29 | 0,56 | 0,64 |
| g:Anaerosporebacter | 0,69 | -0,53 | 0,60 | 0,67 |
| g:Bilophila | 0,79 | -0,33 | 0,67 | 0,73 |
| g:Enterococcus | 1,46 | 0,54 | 0,67 | 0,73 |
| g:Lactobacillus | 1,17 | 0,23 | 0,73 | 0,78 |
| g:Alloprevotella | 1,17 | 0,23 | 0,75 | 0,79 |
| g:Olsenella | 1,13 | 0,17 | 0,84 | 0,86 |
| g:Senegalimassilia | 0,86 | -0,21 | 0,85 | 0,86 |
| g:Oscillibacter | 1,01 | 0,01 | 0,99 | 0,99 |
| **Related to Figure S1** |  |  |  |  |
| **Genus** | **Fold change** | **Log2FC** | **p value** | **Adj p value** |
| g:Akkermansia | 0,0007 | -10,44 | 5,67E-35 | 4,08E-33 |
| g:Bacillus | 83,30 | 6,38 | 4,04E-15 | 1,45E-13 |
| g:Acetatifactor | 0,20 | -2,30 | 9,26E-12 | 2,22E-10 |
| g:Lactobacillus | 32,59 | 5,03 | 1,43E-11 | 2,46E-10 |
| g:Parasutterella | 15,59 | 3,96 | 1,71E-11 | 2,46E-10 |
| g:Clostridium_IV | 0,17 | -2,53 | 3,98E-11 | 4,78E-10 |
| g:Nocardioides | 52,24 | 5,71 | 9,07E-11 | 9,33E-10 |
| g:Clostridium_XVIII | 0,09 | -3,54 | 3,02E-09 | 2,72E-08 |
| g:Clostridium_XIVa | 2,25 | 1,17 | 1,36E-08 | 1,09E-07 |
| g:Sporobacter | 0,21 | -2,22 | 1,77E-08 | 1,28E-07 |
| g:Brevibacillus | 22,94 | 4,52 | 6,24E-08 | 4,08E-07 |
| g:Eubacterium | 4,77 | 2,25 | 9,72E-08 | 5,83E-07 |
| g:Parabacteroides | 4,21 | 2,07 | 4,94E-07 | 2,74E-06 |
| g:Porphyromonas | 3,77 | 1,92 | 5,48E-07 | 2,82E-06 |
| g:Barnesiella | 3,31 | 1,73 | 1,09E-06 | 5,25E-06 |
| g:Enterococcus | 26,51 | 4,73 | 1,43E-06 | 6,42E-06 |
| g:Sphingopyxis | 21,41 | 4,42 | 1,72E-06 | 7,28E-06 |
| g:Anaerovorax | 5,93 | 2,57 | 2,10E-05 | 8,41E-05 |
| g:Haloplasma | 3,38 | 1,76 | 9,88E-05 | 0,0004 |
| g:Arthrobacter | 9,18 | 3,20 | 0,0003 | 0,001 |
| g:Rhizobium | 8,90 | 3,15 | 0,0003 | 0,001 |
| g:Vampirovibrio | 6,25 | 2,64 | 0,0003 | 0,001 |
| g:Allobaculum | 4,86 | 2,28 | 0,0004 | 0,001 |
| g:Neorhizobium | 8,18 | 3,03 | 0,001 | 0,002 |
| g:Paenibacillus | 7,85 | 2,97 | 0,001 | 0,002 |
| g:Romboutsia | 4,91 | 2,29 | 0,001 | 0,002 |
| g:Clostridium_sensu_stricto | 12,44 | 3,64 | 0,001 | 0,002 |
| **g:Anaerotruncus** | **0,48** | **-1,07** | **0,001** | **0,004** |
| **g:Butyricicoccus** | **0,22** | **-2,21** | **0,001** | **0,004** |
| **g:Staphylococcus** | **6,33** | **2,66** | **0,002** | **0,01** |
| **g:Catenisphaera** | **7,26** | **2,86** | **0,002** | **0,01** |
| **g:Catabacter** | **0,47** | **-1,10** | **0,003** | **0,01** |
| **g:Papillibacter** | **0,43** | **-1,23** | **0,003** | **0,01** |
| **g:Coprobacillus** | **0,06** | **-4,00** | **0,01** | **0,01** |
| **g:Dorea** | **0,45** | **-1,16** | **0,01** | **0,02** |
| **g:Lutispora** | **2,96** | **1,56** | **0,02** | **0,04** |
| **g:Shuttleworthia** | **0,17** | **-2,54** | **0,02** | **0,04** |
| g:Alloprevotella | 0,51 | -0,98 | 0,03 | 0,05 |
| g:Pseudoflavonifractor | 0,63 | -0,67 | 0,03 | 0,05 |
| g:Hydrogenoanaerobacterium | 2,49 | 1,32 | 0,03 | 0,06 |
| g:Clostridium XIVb | 0,74 | -0,44 | 0,08 | 0,14 |
| g:Intestinimonas | 1,79 | 0,84 | 0,08 | 0,14 |
| g:Turicibacter | 3,32 | 1,73 | 0,08 | 0,14 |
| g:Prevotella | 3,25 | 1,70 | 0,08 | 0,14 |
| g:Erysipelotrichaceae incertae sedis | 3,62 | 1,86 | 0,12 | 0,19 |
| g:Roseburia | 0,54 | -0,88 | 0,14 | 0,21 |
| g:Christensenella | 2,02 | 1,01 | 0,18 | 0,27 |
| g:Desulfovibrio | 0,80 | -0,33 | 0,23 | 0,34 |
| g:Lachnospiracea incertae sedis | 1,35 | 0,43 | 0,22 | 0,34 |
| g:Murimonas | 1,78 | 0,83 | 0,23 | 0,34 |
| g:Anaerosporeobacter | 0,59 | -0,76 | 0,25 | 0,35 |
| g:Odoribacter | 1,55 | 0,63 | 0,31 | 0,42 |
| g:Enterorhabdus | 1,47 | 0,56 | 0,33 | 0,45 |
| g:Blautia | 0,61 | -0,71 | 0,37 | 0,47 |
| g:Butyrivibrio | 0,70 | -0,51 | 0,37 | 0,47 |
| g:Oscillibacter | 1,23 | 0,30 | 0,37 | 0,47 |
| g:Streptococcus | 0,69 | -0,54 | 0,36 | 0,47 |
| g:Olsenella | 0,53 | -0,91 | 0,46 | 0,58 |
| g:Bilophila | 1,15 | 0,20 | 0,53 | 0,64 |
| g:Coproccoccus | 1,32 | 0,40 | 0,54 | 0,64 |
| g:Aestuariaispira | 0,73 | -0,46 | 0,57 | 0,67 |
| g:Bacteroides | 0,84 | -0,26 | 0,58 | 0,67 |
| g:Ruminococcus2 | 2,00 | 1,00 | 0,58 | 0,67 |
| g:Saccharofermentans | 1,16 | 0,22 | 0,64 | 0,72 |
| g:Gracilibacter | 0,88 | -0,18 | 0,75 | 0,83 |
| g:Alkaliphilus | 0,84 | -0,25 | 0,77 | 0,84 |
| g:Alistipes | 0,94 | -0,09 | 0,85 | 0,89 |
| g:Flavonifractor | 1,04 | 0,06 | 0,86 | 0,89 |
| g:Gemella | 0,90 | -0,15 | 0,88 | 0,89 |
| g:Ruminococcus | 1,05 | 0,07 | 0,87 | 0,89 |
| g:Senegalimassilia | 0,90 | -0,15 | 0,87 | 0,89 |
| g:Clostridium III | 1,01 | 0,01 | 0,96 | 0,96 |

**Table S4:** Lipid species altered by McB: Reported p-values compared to the WD_REF_ are obtained by mixed-effects analysis with multiple comparisons and Dunnett’s post hoc.

| Lipid | Adjusted p-value between<br>WD<sub>REF</sub> and LFD | Adjusted p-value between<br>WD<sub>REF</sub> and WD<sub>McB</sub> |
| --- | --- | --- |
| **Upregulated by McB:** |  |  |
| lysoPC 17:1 | 0,7072 | 0,0022 |
| PC 33:1 B | 0,5237 | <0,0001 |
| PC 33:5 D | 0,0001 | <0,0001 |
| PC 38:4 C | <0,0001 | <0,0001 |
| PC 39:6 | 0,0021 | <0,0001 |
| PE 38:5 | 0,0004 | 0,0081 |
| plasmeyl-PC 36:3 | 0,0002 | 0,0004 |
| **Downregulated by McB:** |  |  |
| lysoPC 16:1 | 0,9977 | 0,0023 |
| lysoPC 18:1 | 0,4575 | 0,0065 |
| lysoPC 20:1 | 0,0003 | 0,0001 |
| lysoPC 20:2 | 0,0027 | 0,0028 |
| lysoPC 20:3 | 0,0019 | 0,0005 |
| lysoPC 20:4 A | 0,9830 | 0,0018 |
| PC 32:1 | 0,0004 | 0,0012 |
| PC 33:5 C | 0,0933 | <0,0001 |
| PC 34:3 | 0,4656 | 0,0001 |
| PC 35:4 | <0,0001 | 0,0003 |
| PC 36:3 A | 0,0003 | 0,0002 |
| PC 36:3 B | <0,0001 | <0,0001 |
| PC 36:5 B | 0,9151 | <0,0001 |
| PC 37:4 A | 0,0009 | 0,0085 |
| PC 38:4 A | 0,0001 | 0,0004 |
| PC 38:4 B | <0,0001 | <0,0001 |
| PC 38:5 A | 0,2436 | <0,0001 |
| PC 38:6 B | 0,0006 | <0,0001 |
| PC 39:7 | 0,0022 | 0,0033 |
| PC 40:7 B | 0,0006 | 0,0001 |

**Table S5:** Deseq analysis from 21+5 week experiment. Results from deseq analysis of bacterial genera of the 21+5 week experiment featuring genus name, Log2 fold change, comparing WD_McB_ to WD_REF_. Unadjusted p-values as well as p-values adjusted by Benjamini-Hochberg method are reported and statistically significantly affected genera (Adj. p < 0.05) are marked in bold.

| Related to Figure 6 |  |  |  |  |
| --- | --- | --- | --- | --- |
| Genus | Fold change | Log2FC | p value | Adj p value |
| g:Parasutterella | 2066,66 | 11,013 | 1,4E-38 | 9,3E-37 |
| g:Parabacteroides | 94,91 | 6,568 | 7,6E-33 | 2,6E-31 |
| g:Brevibacillus | 138,42 | 7,113 | 9,1E-18 | 2,1E-16 |
| g:Pseudoflavonifractor | 0,09 | -3,418 | 3,4E-14 | 5,9E-13 |
| g:Odoribacter | 19,74 | 4,303 | 1,9E-13 | 2,6E-12 |
| g:Desulfovibrio | 0,18 | -2,501 | 4,8E-09 | 5,5E-08 |
| g:Haloplasma | 13,78 | 3,785 | 7,0E-09 | 6,9E-08 |
| g:Dorea | 0,11 | -3,168 | 1,3E-08 | 9,9E-08 |
| g:Flavonifractor | 0,11 | -3,153 | 1,3E-08 | 9,9E-08 |
| g:Acetatifactor | 0,16 | -2,621 | 2,3E-07 | 1,6E-06 |
| g:Rhodococcus | 40,07 | 5,325 | 3,5E-07 | 2,2E-06 |
| g:Romboutsia | 0,04 | -4,556 | 2,9E-06 | 1,7E-05 |
| g:Alistipes | 3,95 | 1,982 | 4,3E-06 | 2,3E-05 |
| g:Porphyromonas | 8,17 | 3,031 | 1,5E-05 | 7,4E-05 |
| g:Shuttleworthia | 0,09 | -3,499 | 1,6E-05 | 7,4E-05 |
| g:Allobaculum | 2,74 | 1,454 | 1,0E-04 | 4,3E-04 |
| g:Barnesiella | 2,88 | 1,524 | 1,1E-04 | 4,7E-04 |
| g:Anaerotruncus | 0,24 | -2,039 | 2,0E-04 | 7,5E-04 |
| g:Akkermansia | 8,07 | 3,013 | 3,5E-04 | 1,3E-03 |
| g:Mucispirillum | 0,14 | -2,789 | 4,8E-04 | 1,7E-03 |
| g:Cohnella | 15,91 | 3,992 | 9,6E-04 | 3,1E-03 |
| g:Clostridium_IV | 0,51 | -0,966 | 1,2E-03 | 3,6E-03 |
| g:Clostridium_XIVa | 0,54 | -0,893 | 1,3E-03 | 3,8E-03 |
| g:Eubacterium | 4,77 | 2,253 | 1,6E-03 | 4,5E-03 |
| g:Papillibacter | 0,27 | -1,874 | 2,1E-03 | 5,8E-03 |
| g:Coproacter | 13,09 | 3,710 | 3,0E-03 | 7,8E-03 |
| g:Oscillibacter | 0,38 | -1,415 | 3,2E-03 | 8,2E-03 |
| g:Bifidobacterium | 7,60 | 2,926 | 4,4E-03 | 1,1E-02 |
| g:Sporobacter | 0,39 | -1,364 | 1,0E-02 | 2,5E-02 |
| **g:Clostridium XVIII** | **0,36** | **-1,481** | **1,2E-02** | **2,8E-02** |
| **g:Ruminococcus** | **0,44** | **-1,178** | **1,6E-02** | **3,6E-02** |
| **g:Catabacter** | **0,59** | **-0,768** | **2,1E-02** | **4,6E-02** |
| **g:Christensenella** | **4,25** | **2,086** | **2,3E-02** | **4,9E-02** |
| g:Clostridium XIVb | 2,63 | 1,395 | 0,025 | 0,050 |
| g:Hydrogenoanaerobacterium | 0,25 | -2,005 | 0,029 | 0,058 |
| g:Streptococcus | 0,38 | -1,406 | 0,033 | 0,064 |
| g:Butyrivibrio | 1,64 | 0,715 | 0,041 | 0,077 |
| g:Lactobacillus | 2,02 | 1,018 | 0,046 | 0,083 |
| g:Olsenella | 2,39 | 1,260 | 0,069 | 0,123 |
| g:Butyricicoccus | 0,31 | -1,678 | 0,079 | 0,137 |
| g:Blautia | 0,47 | -1,096 | 0,113 | 0,182 |
| g:Pedobacter | 3,78 | 1,918 | 0,112 | 0,182 |
| g:Roseburia | 0,52 | -0,954 | 0,111 | 0,182 |
| g:Murimonas | 0,28 | -1,855 | 0,129 | 0,203 |
| g:Fodinicurvata | 3,54 | 1,822 | 0,140 | 0,214 |
| g:Holdemania | 1,96 | 0,970 | 0,150 | 0,225 |
| g:Clostridium III | 1,42 | 0,506 | 0,184 | 0,264 |
| g:Intestinimonas | 0,54 | -0,877 | 0,183 | 0,264 |
| g:Anaerofilum | 0,44 | -1,168 | 0,191 | 0,269 |
| g:Anaerosolibacter | 0,32 | -1,630 | 0,208 | 0,287 |
| g:Coproccoccus | 2,06 | 1,042 | 0,242 | 0,327 |
| g:Erysipelotrichaceae incertae sedis | 0,44 | -1,194 | 0,323 | 0,428 |
| g:Bilophila | 0,68 | -0,546 | 0,329 | 0,428 |
| g:Anaerovorax | 1,79 | 0,843 | 0,352 | 0,449 |
| g:Enterococcus | 2,38 | 1,251 | 0,366 | 0,459 |
| g:Staphylococcus | 0,61 | -0,707 | 0,379 | 0,468 |
| g:Escherichia/Shigella | 0,42 | -1,250 | 0,411 | 0,498 |
| g:Bacteroides | 0,82 | -0,293 | 0,420 | 0,500 |
| g:Lachnospiraceae incertae sedis | 0,75 | -0,408 | 0,469 | 0,548 |
| g:Lutispora | 1,83 | 0,871 | 0,493 | 0,567 |
| g:Alkaliphilus | 1,55 | 0,632 | 0,521 | 0,589 |
| g:Clostridium sensu stricto | 1,60 | 0,680 | 0,562 | 0,625 |
| g:Enterorhabdus | 1,27 | 0,346 | 0,573 | 0,625 |
| no match | 2,30 | 1,202 | 0,579 | 0,625 |
| g:Gracilibacter | 1,23 | 0,304 | 0,591 | 0,627 |
| g:Saccharofermentans | 1,12 | 0,167 | 0,773 | 0,808 |
| g:Anaerosporeobacter | 0,97 | -0,047 | 0,938 | 0,952 |
| g:Tissierella | 1,06 | 0,078 | 0,938 | 0,952 |
| g:Spiroplasma | 1,03 | 0,038 | 0,969 | 0,969 |

## References

1. Honda, K. & Littman, D. R. The microbiota in adaptive immune homeostasis and disease. Nature 535, 75–84 (2016).

2. Koh, A., De Vadder, F., Kovatcheva-Datchary, P. & Bäckhed, F. From dietary fiber to host physiology: Short-chain fatty acids as key bacterial metabolites. Cell 165, 1332–1345 (2016).

3. Lynch, S. V. & Pedersen, O. The Human Intestinal Microbiome in Health and Disease. N. Engl. J. Med. 375, 2369–2379 (2016).

4. Cani, P. D. Human gut microbiome: Hopes, threats and promises. Gut 1–10 (2018). doi:10.1136/gutjnl-2018-316723

5. Seregin, S. S. et al. NLRP6 Protects Il10−/−Mice from Colitis by Limiting Colonization of Akkermansia muciniphila. Cell Rep. 19, 733–745 (2017).

6. Shono, Y. et al. Increased GVHD-related mortality with broad-spectrum antibiotic use after allogeneic hematopoietic stem cell transplantation in human patients and mice. Sci. Transl. Med. 8, 339ra71 (2016).

7. Kovatcheva-Datchary, P. et al. Dietary Fiber-Induced Improvement in Glucose Metabolism Is Associated with Increased Abundance of Prevotella. Cell Metab. 1–12 (2015). doi:10.1016/j.cmet.2015.10.001

8. Nathalie Rolhion, Benoit Chassaing, Marie-Anne Nahori, Jana de Bodt, Alexandra Moura, Marc Lecuit, Olivier Dussurget, Marion Bérard, Massimo Marzorati, Hannah Fehlner-Peach, Dan R. Littman, Andrew T. Gewirtz, Tom Van de Wiele, Pascale CossartNathalie Rolh, P. C. Specific targeting of intestinal Prevotella copri by a Listeria monocytogenes bacteriocin. BIORxIV (2019). doi:https://doi.org/10.1101/680801

9. Pedersen, H. K. et al. Human gut microbes impact host serum metabolome and insulin sensitivity. Nature 535, 376–81 (2016).

10. Plovier, H. et al. A purified membrane protein from Akkermansia muciniphila or the pasteurized bacterium improves metabolism in obese and diabetic mice. Nat. Med. 23, 107– 113 (2017).

11. Verma, R. et al. Cell surface polysaccharides of Bifidobacterium bifidum induce the generation of Foxp3+ regulatory T cells. Sci. Immunol. 3, (2018).

12. Yang, B.-H. et al. Foxp3+ T cells expressing RORγt represent a stable regulatory T-cell effector lineage with enhanced suppressive capacity during intestinal inflammation. Mucosal Immunol. 205, 1381–1393 (2015).

13. Cavallari, J. F., Denou, E., Foley, K. P., Khan, W. I. & Schertzer, J. D. Different Th17 immunity in gut, liver, and adipose tissues during obesity: the role of diet, genetics, and microbes. Gut Microbes 7, 82–89 (2016).

14. Garidou, L. et al. The Gut Microbiota Regulates Intestinal CD4 T Cells Expressing RORγt and Controls Metabolic Disease. Cell Metab. 22, 100–112 (2015).

15. Hong, C. et al. Gut-Specific Delivery of T-Helper 17 Cells Reduces Obesity and Insulin Resistance in Mice. Gastroenterology 152, 1998–2010 (2017).

16. Makula, R. A. Phospholipid composition of methane-utilizing bacteria. J. Bacteriol. 134, 771–777 (1978).

17. Kleiveland, C. R. et al. The noncommensal bacterium Methylococcus capsulatus (Bath) ameliorates dextran sulfate (Sodium Salt)-Induced Ulcerative Colitis by influencing mechanisms essential for maintenance of the colonic barrier function. Appl. Environ. Microbiol. 79, 48–56 (2013).

18. Damgaard, M. T. F. et al. Age-dependent alterations of glucose clearance and homeostasis are temporally separated and modulated by dietary fat. J. Nutr. Biochem. 54, (2018).

19. Kleiner, D. E. et al. Design and validation of a histological scoring system for nonalcoholic fatty liver disease. Hepatology 41, 1313–1321 (2005).

20. Schneider, C. A., Rasband, W. S. & Eliceiri, K. W. NIH Image to ImageJ: 25 years of image analysis. Nat. Methods 9, 671–675 (2012).

21. Ruifrok, A. C. & Johnston, D. A. Quantification of histochemical staining by color deconvolution. Anal. Quant. Cytol. Histol. 23, 291–9 (2001).

22. Ehmann, D. et al. Paneth cell α-defensins HD-5 and HD-6 display differential degradation into active antimicrobial fragments. Proc. Natl. Acad. Sci. 201817376 (2019). doi:10.1073/pnas.1817376116

23. Larsen, I. S. et al. Human Paneth cell α-defensin-5 treatment reverses dyslipidemia and improves glucoregulatory capacity in diet-induced obese mice. Am. J. Physiol. Metab. 317, E42–E52 (2019).

24. Love, M. I., Huber, W. & Anders, S. Moderated estimation of fold change and dispersion for RNA-seq data with DESeq2. Genome Biol. 15, 550 (2014).

25. Luda, K. M. et al. IRF8 Transcription-Factor-Dependent Classical Dendritic Cells Are Essential for Intestinal T Cell Homeostasis. Immunity 44, 860–874 (2016).

26. Bisanz, J. E., Upadhyay, V., Turnbaugh, J. A., Ly, K. & Turnbaugh, P. J. Meta-Analysis Reveals Reproducible Gut Microbiome Alterations in Response to a High-Fat Diet. Cell Host Microbe 26, 265–272.e4 (2019).

27. Ju, T., Kong, J. Y., Stothard, P. & Willing, B. P. Defining the role of Parasutterella, a previously uncharacterized member of the core gut microbiota. ISME J. 13, 1520–1534 (2019).

28. Wang, K. et al. Parabacteroides distasonis Alleviates Obesity and Metabolic Dysfunctions via Production of Succinate and Secondary Bile Acids. Cell Rep. 26, 222–235.e5 (2019).

29. Ottosson, F. et al. Connection Between BMI-Related Plasma Metabolite Profile and Gut Microbiota. J. Clin. Endocrinol. Metab. 103, 1491–1501 (2018).

30. Petersen, C. et al. T cell-mediated regulation of the microbiota protects against obesity. Science 365, eaat9351 (2019).

31. Luck, H. et al. Gut-associated IgA+ immune cells regulate obesity-related insulin resistance. Nat. Commun. 10, 3650 (2019).

32. Macfarlane, G. T. & Macfarlane, S. Bacteria, colonic fermentation, and gastrointestinal health. J. AOAC Int. 95, 50–60 (2012).

33. Tremaroli, V. & Bäckhed, F. Functional interactions between the gut microbiota and host metabolism. Nature 489, 242–249 (2012).

34. Cummings, J. H., Pomare, E. W., Branch, W. J., Naylor, C. P. & Macfarlane, G. T. Short chain fatty acids in human large intestine, portal, hepatic and venous blood. Gut 28, 1221–7 (1987).

35. Mowat, A. M. & Agace, W. W. Regional specialization within the intestinal immune system. Nat. Rev. Immunol. (2014). doi:10.1038/nri3738

36. Xu, M. et al. c-MAF-dependent regulatory T cells mediate immunological tolerance to a gut pathobiont. Nature (2018). doi:10.1038/nature25500

37. Damgaard, M. T. F. et al. Age-dependent alterations of glucose clearance and homeostasis are temporally separated and modulated by dietary fat. J. Nutr. Biochem. 54, 66–76 (2018).

38. Friedman, S. L., Neuschwander-Tetri, B. A., Rinella, M. & Sanyal, A. J. Mechanisms of NAFLD development and therapeutic strategies. Nat. Med. 24, 1 (2018).

39. Giles, D. A. et al. Thermoneutral housing exacerbates nonalcoholic fatty liver disease in mice and allows for sex-independent disease modeling. Nat. Med. 23, 829–838 (2017).

40. Kindt, A. et al. The gut microbiota promotes hepatic fatty acid desaturation and elongation in mice. Nat. Commun. 9, (2018).

41. Singh, V. et al. Microbiota-Dependent Hepatic Lipogenesis Mediated by Stearoyl CoA Desaturase 1 (SCD1) Promotes Metabolic Syndrome in TLR5-Deficient Mice. Cell Metab. 22, 983–996 (2015).

42. Forouhi, N. G. et al. Differences in the prospective association between individual plasma phospholipid saturated fatty acids and incident type 2 diabetes: the EPIC-InterAct case-cohort study. *lancet*. Diabetes Endocrinol. 2, 810–8 (2014).

43. Pagadala, M., Kasumov, T., McCullough, A. J., Zein, N. N. & Kirwan, J. P. Role of ceramides in nonalcoholic fatty liver disease. Trends Endocrinol. Metab. 23, 365–71 (2012).

44. Ju, C. & Tacke, F. Hepatic macrophages in homeostasis and liver diseases: From pathogenesis to novel therapeutic strategies. Cell. Mol. Immunol. 13, 316–327 (2016).

45. Ganeshan, K. & Chawla, A. Warming the mouse to model human diseases. Nat. Publ. Gr. 13, 458–465 (2017).

46. Tian, X. Y. et al. Thermoneutral Housing Accelerates Metabolic Inflammation to Potentiate Atherosclerosis but Not Insulin Resistance_Suppl… Cell Metab. 23, 165–178 (2016).

47. Lathrop, S. K. et al. Peripheral education of the immune system by colonic commensal microbiota. Nature 478, 250–4 (2011).

48. Xiao, L. et al. High-fat feeding rather than obesity drives taxonomical and functional changes in the gut microbiota in mice. Microbiome 5, 1–12 (2017).

49. Zhang, C. et al. Structural resilience of the gut microbiota in adult mice under high-fat dietary perturbations. ISME J. 6, 1848–1857 (2012).

50. Desai, M. S. et al. A dietary fiber-deprived gut microbiota degrades the colonic mucus barrier and enhances pathogen susceptibility HHS Public Access Graphical abstract. Cell 167, 1339–1353 (2016).

51. Dalby, M. J., Ross, A. W., Walker, A. W. & Morgan, P. J. Dietary Uncoupling of Gut Microbiota and Energy Harvesting from Obesity and Glucose Tolerance in Mice. Cell Rep. 21, 1521–1533 (2017).

52. Lennon, G. et al. Correlations between colonic crypt mucin chemotype, inflammatory grade and Desulfovibrio species in ulcerative colitis. Color. Dis. 16, 161–169 (2014).

53. Dawson, P. A. et al. Reduced mucin sulfonation and impaired intestinal barrier function in the hyposulfataemic NaS1 null mouse. Gut 58, 910–919 (2009).

54. Shan, M. et al. Mucus enhances gut homeostasis and oral tolerance by delivering immunoregulatory signals. Science (80-.). 342, 447–453 (2013).

55. Persson, E. K., Scott, C. L., Mowat, A. M. & Agace, W. W. Dendritic cell subsets in the intestinal lamina propria: Ontogeny and function. Eur. J. Immunol. 43, 3098–3107 (2013).

56. Christoffersen, T. E. et al. Effects of the non-commensal Methylococcus capsulatus Bath on mammalian immune cells. Mol. Immunol. 66, 107–116 (2015).

57. Smith, P. M. et al. The microbial metabolites, short-chain fatty acids, regulate colonic Treg cell homeostasis. Science 341, 569–73 (2013).

58. Karlmark, K. R. et al. Hepatic recruitment of the inflammatory Gr1+ monocyte subset upon liver injury promotes hepatic fibrosis. Hepatology 50, 261–74 (2009).

59. Dal-Secco, D. et al. A dynamic spectrum of monocytes arising from the in situ reprogramming of CCR2+ monocytes at a site of sterile injury. J. Exp. Med. 212, 447–56 (2015).

60. Ramachandran, P. et al. Differential Ly-6C expression identifies the recruited macrophage phenotype, which orchestrates the regression of murine liver fibrosis. Proc. Natl. Acad. Sci. U. S. A. 109, E3186–95 (2012).

61. Moschen, A. R., Tilg, H. & Raine, T. IL-12, IL-23 and IL-17 in IBD: immunobiology and therapeutic targeting. Nat. Rev. Gastroenterol. Hepatol. 16, 185–196 (2019).

62. Bregenzer, N. et al. Increased insulin resistance and beta cell activity in patients with Crohn’s disease. Inflamm. Bowel Dis. 12, 53–6 (2006).

63. Chassaing, B. et al. Lack of soluble fiber drives diet-induced adiposity in mice. Am. J. Physiol. - Gastrointest. Liver Physiol. 309, G528–G541 (2015).

64. Zou, J. et al. Fiber-Mediated Nourishment of Gut Microbiota Protects against Diet-Induced Obesity by Restoring IL-22-Mediated Colonic Health. Cell Host Microbe 23, 41–53.e4 (2018).

65. Stange, E. F. & Schroeder, B. O. Microbiota and mucosal defense in IBD: an update. Expert Rev. Gastroenterol. Hepatol. 00, 1–14 (2019).

66. Yang, L. et al. Amelioration of high fat diet induced liver lipogenesis and hepatic steatosis by interleukin-22. J. Hepatol. 53, 339–347 (2010).

67. Miani, M. et al. Gut Microbiota-Stimulated Innate Lymphoid Cells Support β-Defensin 14 Expression in Pancreatic Endocrine Cells, Preventing Autoimmune Diabetes. Cell Metab. 1– 16 (2018). doi:10.1016/j.cmet.2018.06.012

