## Supplementary figures and images for "Lysates of *Methylococcus capsulatus* Bath induce a lean-like microbiota, intestinal FoxP3^+^RORγt^+^IL-17^+^ Tregs and improve metabolism"

### Supplemental Figure 1

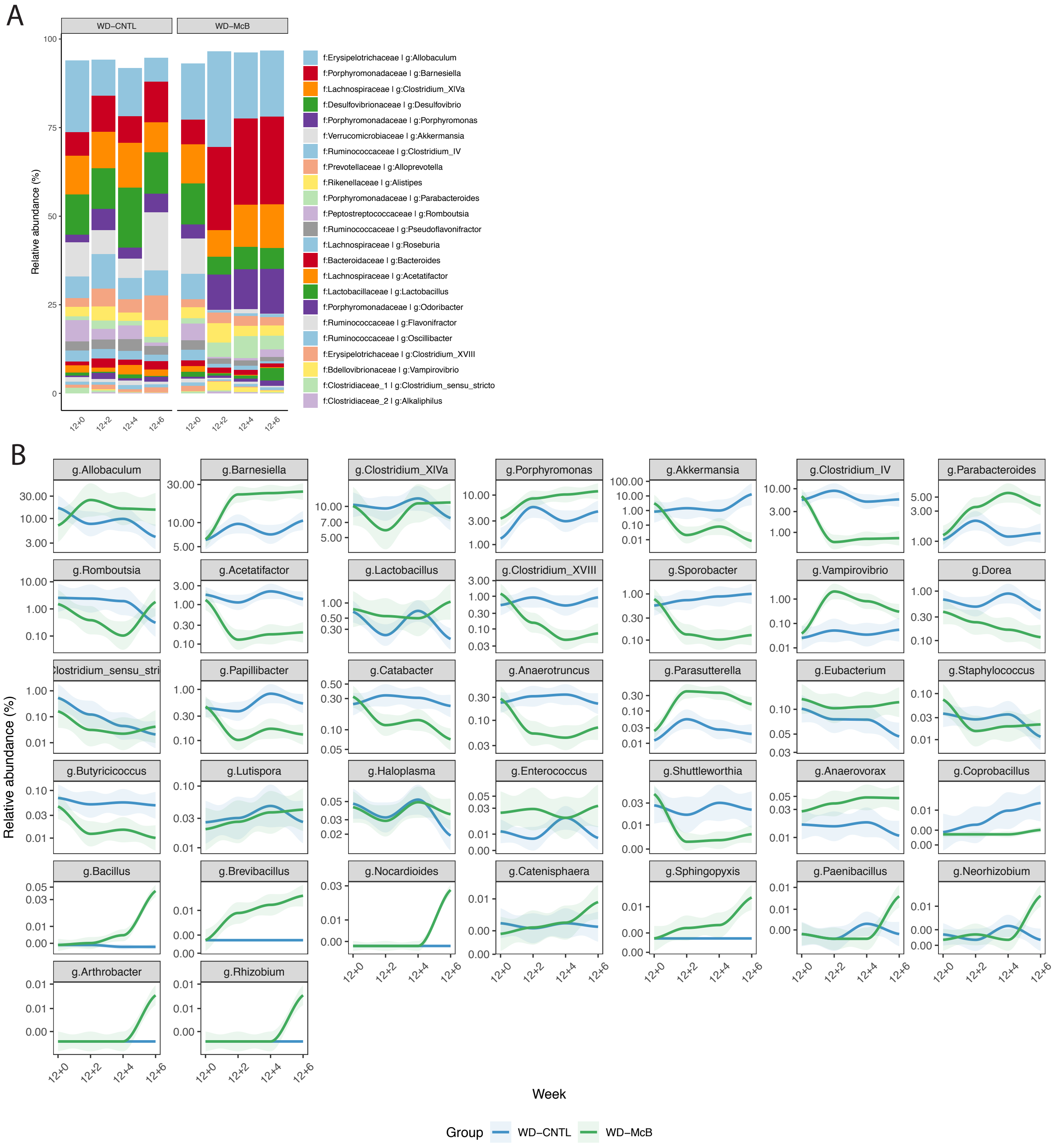

### Supplemental Figure 2

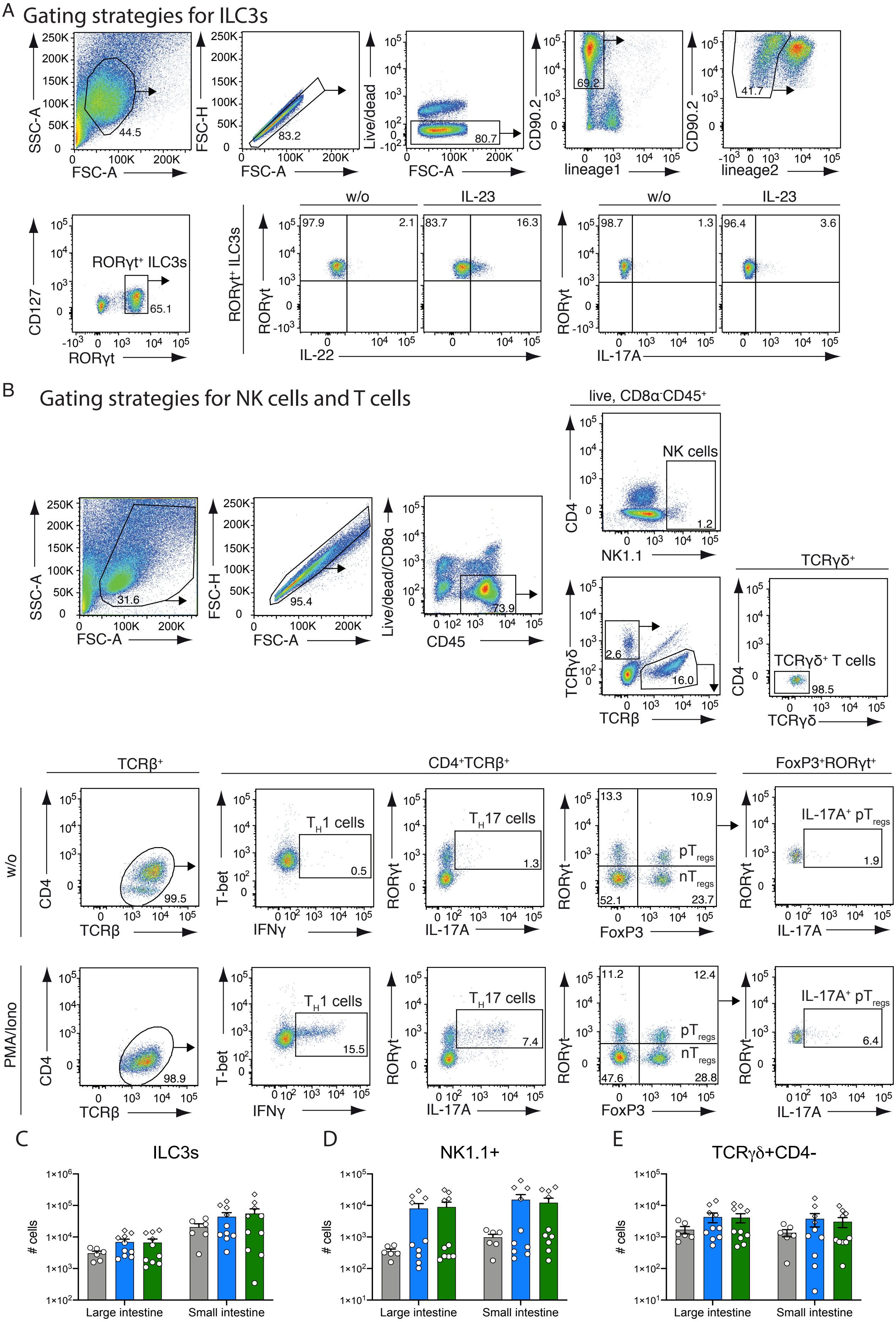

### Supplemental Figure 3

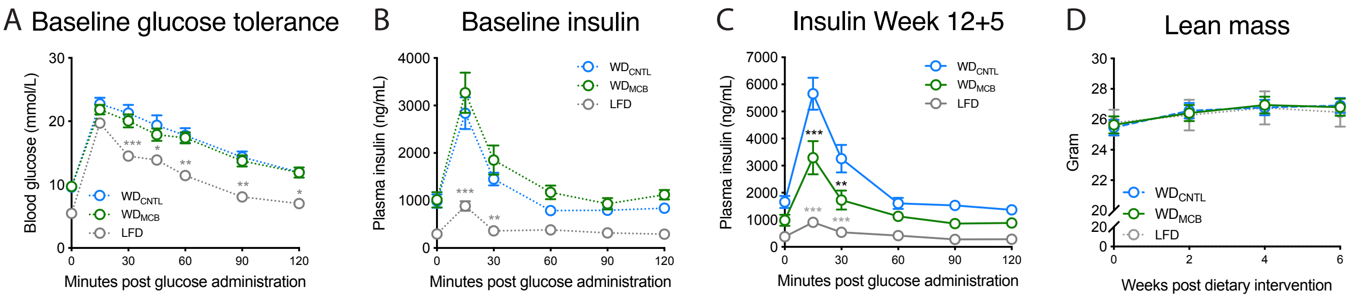

### Supplemental Figure 4

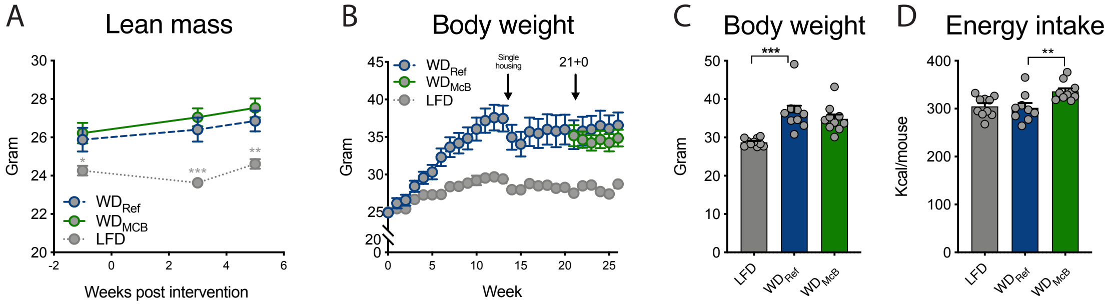

### Supplemental Figure 5

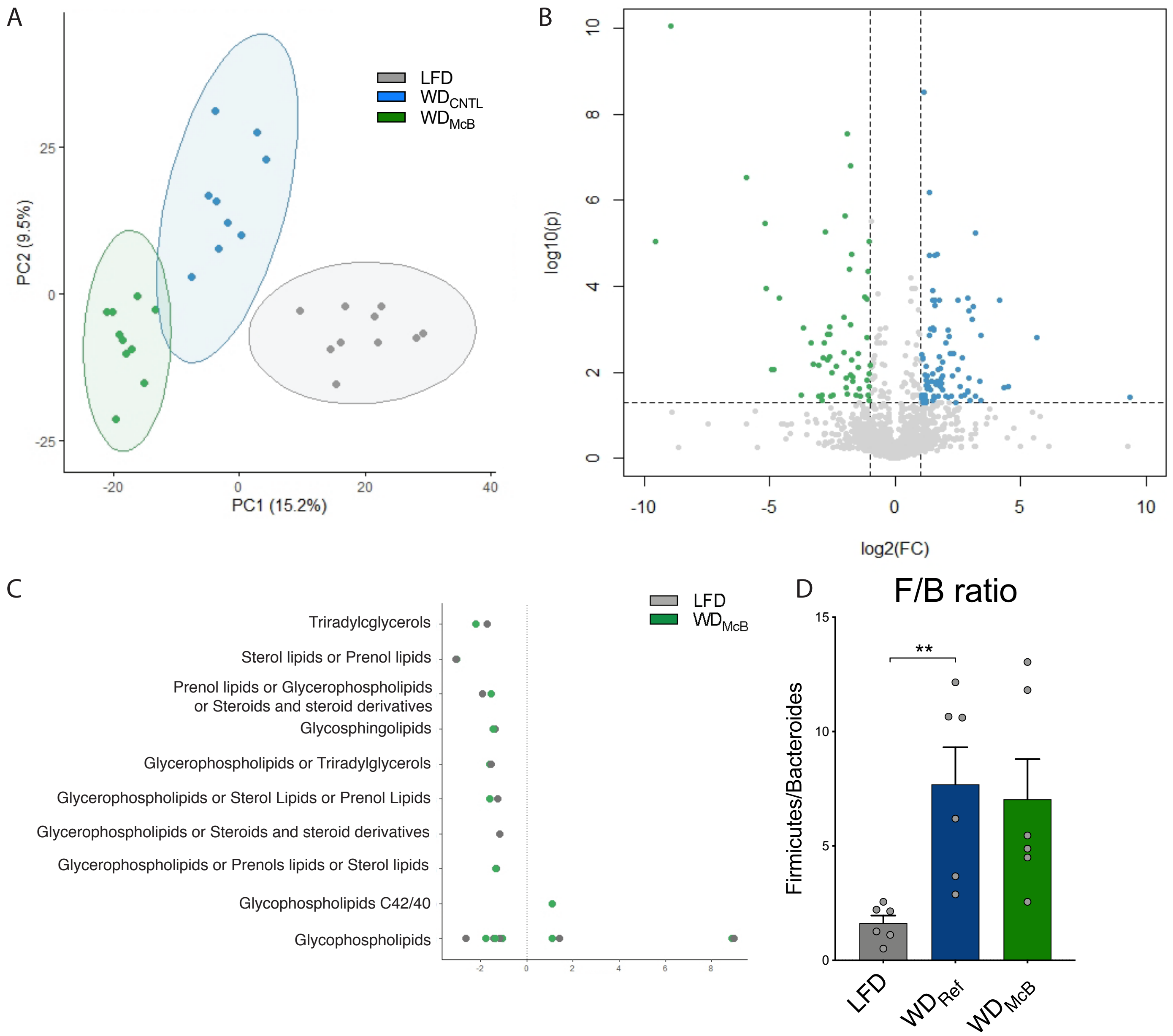
